# Amount of fiction reading correlates with higher connectivity between cortical areas for language and mentalizing

**DOI:** 10.1101/2020.06.08.139923

**Authors:** Franziska Hartung, Roel M. Willems

## Abstract

Behavioral evidence suggests that engaging with fiction is positively correlated with social abilities. The rationale behind this link is that engaging with fiction and fictional characters may offer a ‘training mode’ for mentalizing and empathizing with sentient agents in the real world, analogous to a flight simulator for pilots. In this study, we explored the relationship between reading fiction and mentalizing by looking at brain network dynamics in 57 participants who varied on how much fiction they read in their daily lives. The hypothesis was that if reading fiction indeed trains mentalizing, a task that requires mentalizing –Like immersing in a fictional story and engaging with a protagonist-should elicit differences in brain network dynamics depending on how much people read. More specifically, more frequent readers should show increased connectivity within the theory of mind network (ToM) or between the ToM network and other brain networks. While brain activation was measured with fMRI, participants listened to two literary narratives. We computed time-course correlations between brain regions and compared the correlation values from listening to narratives to listening to an auditory baseline condition. The between-region correlations were then related to individual differences measures including the amount of fiction that participants consume in their daily lives. Our results show that there is a linear relationship between how much people read and the functional connectivity in areas known to be involved in language and mentalizing. This adds neurobiological credibility to the ‘fiction influences mentalizing abilities’ hypothesis as suggested on the basis of conceptual analysis.

## Introduction

It has been hypothesized since antiquity that people who engage with fiction have better social skills (1–3). A proposed mechanism behind this link is that engaging with fictional narratives offers a ‘training mode’ for social cognition (4–10). The idea is that just as flight simulators train the skills of pilots, engaging with narratives enables the individual to simulate interacting with social agents, and thereby train the capacity to understand the intentions, feelings, desires, and beliefs of other people.

Researchers have focused most prominently on the effect of fiction on *empathizing* and *mentalizing* abilities (7). Empathizing refers to the capacity to experience feelings from another person’s perspective. Mentalizing refers to the capacity to understand and reason about other individuals’ intentions and beliefs. While empathy and mentalizing are separable psychological constructs with separable neural (11), there is no specific prediction for either system and the hypothesis predicts beneficial effects of fiction reading on social cognition more generally. Hence, we treat empathy and mentalizing as closely related for the purpose of the present study and refer to them together as *social cognition* for the remainder of the paper.

While it is difficult to prove a causal relationship between reading fiction and social cognition abilities, recent behavioral evidence confirms that engaging with fiction (such as written narratives) is positively correlated with better social cognition performance and self-report abilities. For instance, Mar and colleagues (12) found that lifetime fiction exposure was positively correlated with performance on several social cognition measures (see also 13). The correlation of lifetime amount of reading with empathizing skills has been replicated in several large-scale studies, and hence seems to be a robust phenomenon (14,15). In addition to this life-time (‘long-term’) effect, some behavioral evidence suggests that prosocial behavior can be increased immediately after fiction exposure (16), and even a direct improvement of social cognition abilities after only brief exposure to fiction (17–19). However, three recent large-scale replication studies of the original effect showed that brief exposure to fiction in fact does not significantly improve social cognition (14,15,20).

Despite the behavioral evidence for the correlation between fiction reading and social cognition skillset, it is unclear what an underlying neural mechanism that supports such a social training mode would look like. Neuroimaging is a promising tool to investigate the relation between social cognition and fiction reading since we can rely on a well-established set of brain regions which are known to be activated by mentalizing and empathizing tasks (21,22). The brain networks involved in social cognition are the so-called mentalizing or Theory of Mind (ToM) network and the empathy network. Both networks are functionally separable from each other (11) as well as from the neural network involved in the basic aspects of language comprehension, such as semantic and syntactic processing (23). The ToM brain network partially overlaps with the default mode network (DMN) (24), which again overlaps with regions that process language at the level of discourse such as narratives (25,26), also known as the extended language network (27). The DMN is sensitive to coherence, and processes information over time scales of several minutes (25,26). It is involved in episodic thinking, social cognition, self-referential thinking, and daydreaming/ mind wandering which all involve narrative structure of some sort (28). While the investigation of the link between reading and social cognition is fairly new in neuroimaging research, one recent study showed that parts of the mentalizing network were more strongly activated in frequent readers during comprehension of social content in brief excerpts of fiction (13).

In the present study, we investigate a possible underlying neural mechanism behind the link between how regularly people engage with fiction, and social cognition, by investigating connections between brain areas while people listened to literary short stories. While listening and reading stories might train slightly different parts of the brain, higher order areas affected by semantic and social factors should be invariant to input modality (29–32). We aim to provide insight into the neurobiological mechanisms that links social cognition abilities and engagement with narratives. We had a sample of healthy young participants (N=57)Listen to two literary short stories while brain activation was measured with functional Magnetic Resonance Imaging (fMRI). Participants also listened to a reversed speech version of the same stories, which served as a low-level baseline. After the scanning session, participants filled out a questionnaire battery including a brief questionnaire related to their reading habits.

In the analysis we extracted the time courses during listening to the narratives and listening to reversed speech separately for each participant from brain regions spanning the whole cortical sheet. We then computed correlations between all regions and computed the difference in correlation values between listening to narratives versus listening to reversed speech. This difference was correlated with individual differences in fiction reading across participants. Individual differences in brain connectivity are more robust in connected stimulus presentation such as movies or narratives compared to isolated event related stimuli or rest (33).

In our analysis, we targeted the whole cortical sheet in order to not exclude certain regions a priori. We did however have hypotheses about certain networks in which we expected that differences in reading habits would have an influence on connectivity with other regions. First, these are the regions of the mentalizing or ToM network (in the remainder of the paper we will refer to these as ‘mentalizing network’) encompassing the anterior part of the medial prefrontal cortex, the temporo-parietal junctions bilaterally, and the anterior temporal cortex bilaterally (22,34,35). Second, as regions involved in empathizing we define the anterior insulae, and the anterior cingulate cortex (ACC) (11). If there is an influence of fiction reading on social cognition abilities, we expect the density of functional connections of parts of the mentalizing and empathizing brain networks to be positively related to the amount of fiction reading that our participants report in their daily lives.

Since reading is related to language, we also expected that brain regions involved in language comprehension are sensitive to how much people read (36), and might show covariation and connection with the mentalizing regions. Among the regions we expected to be involved were the inferior frontal cortex bilaterally, the left posterior middle temporal gyrus, and the left angular gyrus (37–40). Some of these areas are also known to be involved in emotional language processing and mentalizing (41). One reason to not target these networks directly, but instead employ a whole brain analysis approach, was that a lot of previous research has not investigated mentalizing or language processes at the discourse level (with notable exceptions, see Ferstl, 2010 for overview).Limiting our analysis to a priori defined networks would hence increase the chance of overlooking areas that are not described in previous literature (42).

## Methods and Materials

### Participants

Sixty participants took part in the experiment. Data from three participants were removed from subsequent analysis. One participant had a data acquisition artifact in one of the runs, and two other participants had more than 15% of data corrected in at least one of the runs in the motion detection procedure (see below). All analyses are hence performed on data of 57 participants (31 women, 26 men; mean age 22.7 years; s.d. 3.64,Range 18-35; 9Left-handed by self-report, see Willems et al. (43) for reasons to not exclude left-handed participants). All participants were healthy volunteers with Dutch as their native language, and without a history of neurological or psychiatric illness. The study was approved by an external ethics committee (CMO Committee on research Involving Human Subjects, Arnhem-Nijmegen, Netherlands, protocol number 2001/095), and participants gave informed consent in accordance with the Declaration of Helsinki. Participants were paid for participation.

### Stimuli

#### Literary Stories and baseline

Stimuli were audio recordings of two literary short stories, and a reversed speech baseline of parts of the same stories (Table 1). There were two minimally different versions of both stories and each participant listened to one version of each story. The two different versions only differed in the personal pronouns referring to the protagonist of the story. The original study for which this data was recorded showed that the switching of pronouns has no effect on processing or activated brain regions (44). The reversed speech baseline was created by taking the first half of one and the second half of the other story played backwards, rendering it unintelligible.

**Table 1:** The story stimuli. Both stories were from Dutch literature and were presented as audio recordings to the participants. Each story was presented in two formats, which were different only in the pronouns that refer to the protagonist (‘I’ versus ‘She’).

| Story | Number of words | Duration (min:sec) | Author | Year | Publisher, city |
| --- | --- | --- | --- | --- | --- |
| De Mexicaanse Hond<br>(*The Mexican Dog*) | 1239 | 7:17 /<br>7:23 | Margo Minco | 1990 | Bert Bakker,<br>Amsterdam |
| De Muur (*The Wall*) | 1121 | 7:01 /<br>7:04 | Peter Minten | 2013 | De Geus,<br>Breda |

Natural breaks in the content of the stories (paragraph end) were used to define the cut off, which results in the reversed speech recording being slightly longer than both stories. Since reversed speech is unintelligible, we only used half of each recording for the reversed speech baseline to save time on the scanning protocol. The reasoning for including this low-level baseline was to a) be able to subtract activations linked to low level processes like perception in addition to individual intrinsic connectivity, and b) to have a ‘task’ that does not involve introspective or narrative processing to get a reliable estimate of individual resting state connectivity in cortical networks involved in processing discourse and mentalizing.

The materials were spoken by a female native speaker of Dutch, recorded in a sound-proof studio, and digitized at 48 kHz. Both stories were from established writers in Dutch. They were told from the perspective of the protagonist and describe an event from the protagonist’s life (see Supplementary Materials S1 for a translation). The duration of each story was around 7 minutes (Table 1) and the duration of the reversed speech baseline was 7 minutes 25 seconds.

#### Individual difference measures

After the scanning session, participants answered three questions related to their reading behavior: ‘How many novels did you read last year?’ (free response option), ‘Do you like reading fiction?’ (1-7Likert scale, ‘No – Very much’), ‘How often do you read fiction?’ (1- never, 2- I don’t regularly read fiction, 3 - one time per week, 4 - more than twice per week, 5 - daily). They additionally filled out a Dutch version of the Author Recognition Test (ART), which is a widely used implicit measure of print exposure (45,46). The ART requires participants to select existing author names from a list of 42 names of which 30 are existent fiction authors and 12 are made up names. The score of each participant is computed by subtracting the sum of all incorrect answers from the sum of all correct answers (total score can vary between -12 to 30). We included both the questions regarding current reading behavior and the ART because they target slightly different aspects of reading. The ART measures lifetime print exposure, whereas our questions regarding reading targeted more current habits. Since the hypotheses we tested are agnostic to this aspect, we decided to test for both lifetime exposure and current habits.

In addition, participants filled out the 40 item version of the Empathy Quotient (EQ) questionnaire (47). The EQ questionnaire is a self-report measure of trait empathy consisting of 40 questions testing how sensitive individuals are to their own and others’ emotions and is widely used as a diagnostic tool for social and developmental disorders. Individual scores can range between 0 and 80, and a score of 32 or lower indicates atypical performance. For this study, we used the EQ to check the distribution of EQ in our sample. We also conducted a control analysis to test the sensitivity of our analysis to differences in brain connectivity depending on scores on individual differences measures. People who score higher on the EQ should show a modulation of brain connectivity in brain areas related to social cognition depending on their individual EQ scores if these types of correlations are sensible. This tests whether our fiction reading measures and their modulation of brain connectivity is sensible. Participants also filled out the Need for Cognition (NfC) scale, the Need for Affect (NfA) scale, and the Fantasy Scale (FS) of the Interpersonal Reactivity Index. These three measures target individual differences in reward experienced when engaging with difficult cognitive problems (NfC), intense emotions (NfA), and immersing into narratives and role play (FS). These measures were collected for a related project on enjoyment and aesthetic experiences during reading, the results of which are not analyzed here.

### Procedure

Stimuli were presented to participants through MR-compatible ear plugs which additionally served to reduce the noise from the scanner. All participants could hear the stimuli well, as assessed by a volume test with an excerpt from another audio story spoken by the same speaker. The scanner was collecting images during the volume test (and hence making loud noises). Participants indicated whether volume should be increased or decreased during the volume test, to be able to comfortably listen to the materials. None of the participants asked for the volume to be increased to the maximum level, indicating that they could all hear the stories comfortably. Participants always listened to the narratives first. The order of stories was pseudo-randomized, with one order occurring one time more often in the final data set. After each story, participants filled out a brief survey assessing their engagement with and appreciation of the story. The reversed speech baseline was always presented after participants listened to the narratives and was randomized in order with the other two tasks that are not analyzed here (44). In the two tasks we do not analyse here, participants listened to the same stories two more times while engaging in mental imagery. After the scanning session, participants were asked about their general performance in each task and whether they were familiar with any of the stories before listening to them in the scanner. Participants then filled out the individual differences questionnaires outside of the scanner room and were debriefed.

#### fMRI data acquisition and preprocessing

Images of blood-oxygen level dependent (BOLD) signal were acquired on a 3T Siemens Magnetom Trio scanner (Erlangen, Germany) with a 32-channel head coil. Cushions and tape were used to minimize participants’ head movement. Functional images were acquired using a fast T2*-weighted 3D EPI sequence (48), with high temporal resolution (TR: 880 ms, TE: 28 ms, flip angle: 14 degrees, voxel size: 3.5 × 3.5 × 3.5 mm, 36 slices). High resolution (1 × 1 × 1.25 mm) structural (anatomical) images were acquired using an MP-RAGE T1 GRAPPA sequence. Preprocessing was performed using SPM8 (http://www.fil.ion.ucl.ac.uk/spm) and Matlab 2010b (http://www.mathworks.nl/). After removing the first five volumes to control for T1 equilibration effects, images were realigned to the first image in a run using rigid body registration (‘motion correction’). The mean of the motion-corrected images was then brought into the same space as the individual participants’ anatomical scan. The anatomical and functional scans were spatially normalized to the standard MNI template. Voxel sizes were resampled to 2×2×2 mm per voxel. Finally, all data were spatially smoothed using an isotropic 8 mm full width at half maximum (FWHM) Gaussian kernel. Recent studies show that small head movements can influence correlations between regions’ time courses (49). Therefore, we performed an additional correction for small head movements using the ArtRepair toolbox (http://cibsr.stanford.edu/tools/human-brain-project/artrepair-software.html) (50). Images which showed more than 1.5% signal change from the global signal mean, as well as images in which there was more than 0.5mm motion (compared to the previous image), were replaced by the nearest non-repaired scans (49). This led to replacement of on average 1.01% of images per participant (s.d. 0.55%,Range 0-5.81%, median 0.33%). The mean framewise displacement was 4.39 volumes (median = 0; IQR = 4). This was similar across the three runs (mean run 1 = 3.33; mean run 2 = 2.80; mean run 3 = 7.01; median run 1 = 0; median run 2= 0; median run 3 = 0; IQR run1 = 3.5; IQR run 2 = 2; IQR run 3= 7).

#### fMRI network analysis

To compute correlations between cortical regions, we used the Harvard-Oxford cortical brain atlas (51), as provided in the FSL fMRI data analysis package. This atlas subdivides each hemisphere into 48Regions, which means that there are 96Regions in total (Fig. 1A).

**Figure 1:**
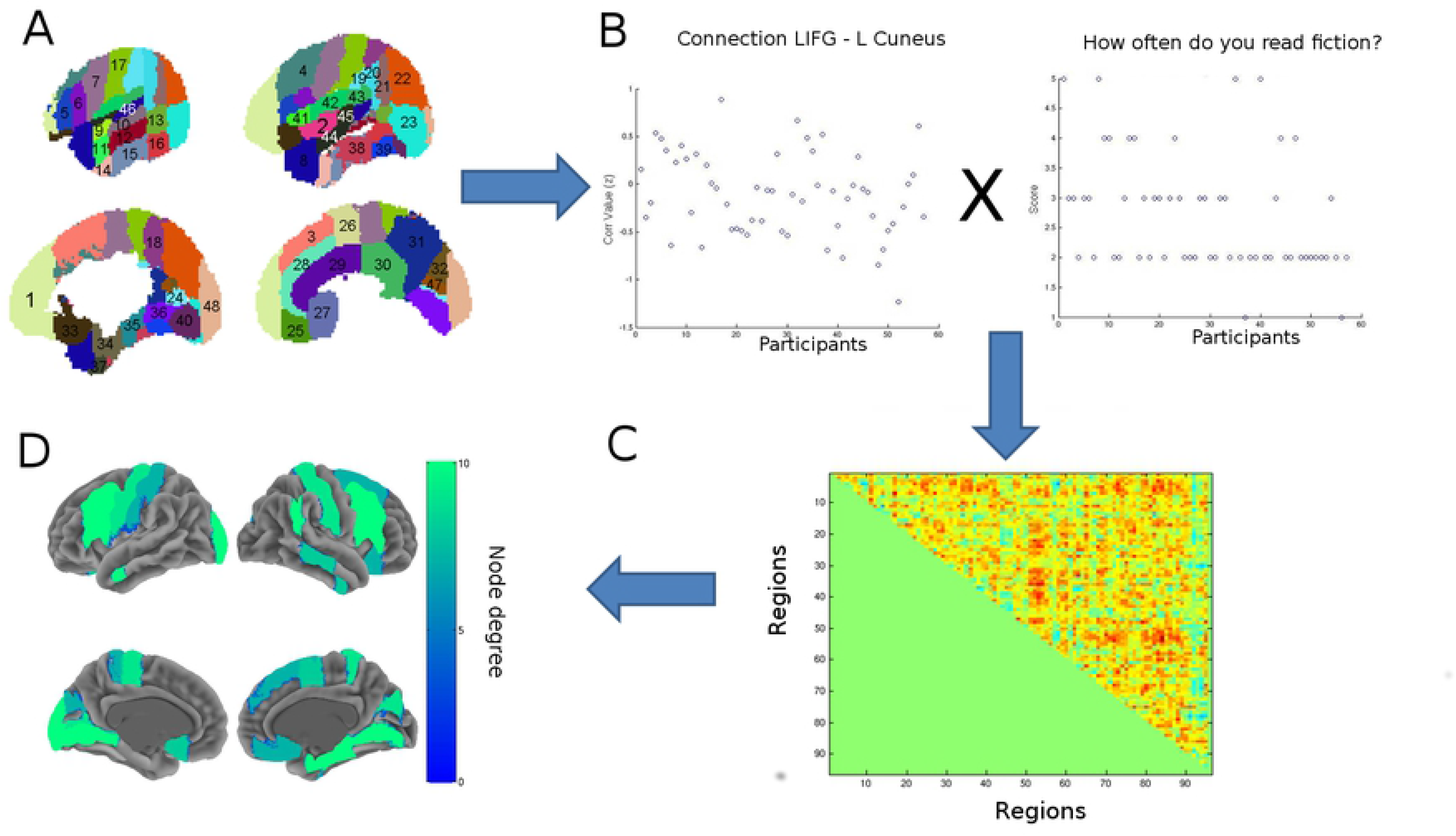
Illustration of analysis steps. A) Time courses were extracted from each of 96 cortical areas in the Harvard-Oxford atlas, covering the whole cortical sheet (for legend see Supplementary Materials S2). B) Spearman correlations were computed between all possible pairs of regions. Correlations during listening to reversed speech baseline were subtracted from correlations during listening to the narratives. The left panel (B1) shows correlation value differences (z-transformed) for one randomly chosen region-region pair for all participants (x-axis, the order of participants is arbitrary). The right panel shows the scores over participants on the post-scanning questionnaire item ‘How often do you read fiction?’ (1-5 scale). For each region-region pair we correlated the between region correlation differences (narrative minus reversed speech) with the scores on the items such as ‘How often do you read fiction’. This gives one correlation value per region-region pair, and the result is illustrated in C) correlation matrix representing the output from step B. The matrix plots per region-region pair the correlation between the correlation differences between those regions’ time courses (B), and the behavioral measure (B). D)Regions with statistically significantly node degree (number of ‘connections’ in C) as compared to the reference distribution based on randomization tests.

We extracted the mean time course per region for each participant. Time courses were detrended (removing linear trend). Next, Spearman’s correlations between each region and all other regions were calculated per run (2 narrative runs and 1Reversed speech). Spearman’s rank correlation was used because it is not sensitive to outliers and does not require normal distribution of the data. Given the distribution of the individual differences data within our sample, Spearman’s rank was the most sensible choice.

The resulting correlation values (3 values for each participant) were transformed to Fisher’s z values. The z-values from the reversed speech runs were subtracted from z-values of the two narrative runs and then averaged (*mean(narrative1-reversed speech,narrative2-reversed speech)*). Subtraction of the reversed speech condition was done to remove any inherent correlations between regions not related to narrative comprehension (which is absent in the reversed speech run), and row-level perceptual processes. Thus, there was one correlation value per region-pair, per participant.

The between-region correlation differences were subsequently correlated with the individual difference scores (see above) using Spearman’s correlation (Fig. 1B). Node degree was calculated for each region from these correlations (Fig. 1C) by counting how many region-region connections had a positive correlation with an individual difference score, at the p<0.01Level (one-sided test) (Fig. 1D). Please note that the connection strengths are not controlled for multiple comparisons. The Family Wise Error Rate is controlled at the revel of node degree per region using a randomization procedure (see below), and the choice of the p<0.01 significance level is arbitrary (see Maris & Oostenveld, (52) for an application of this approach to electrophysiology). Changing this first step p-value does not qualitatively change the overall results. We chose node degree as the dependent variable since it is a well-established basic metric of influence within cortical networks (e.g. (53)). Node degree says nothing about other regions to which a given region is connected. This we illustrate later in the analysis for key regions (see below and Fig. 4). A full overview of connections is given in Supplementary Materials S3. Another reason for taking node degree instead of correlations between regions per se, was the fact that correlation estimates tend to be unstable with sample sizes as employed here. By combining correlation coefficients in node degree, we only look at regions which have higher correlation values for many region-region connections.

A randomization approach was taken to assess which number of connections constitutes a statistically significant effect. The critical value for node degree was established by calculating node degree after randomizing the individual difference scores for each individual difference score separately. This procedure was repeated 2,500 times, and the maximal node degree across regions for each randomization was used to build a reference distribution. The critical value for node degree was the value in the randomization distribution compared to which only 2.5% of randomization produced a more extreme value (corresponding to p<0.05 two-sided, family-wise error rate). In this way, we explicitly tested which regions had a statistically significant number of connections with other regions influenced by a given individual difference score. The analysis showed that 6Regions or more was the maximal critical value across individual difference measures, and we adopted this value to correct for multiple comparisons. Visualizations were created using the Surfplot toolbox written by Aaron Schultz (http://mrtools.mgh.harvard.edu/index.php?title=SurfPlot).

## Results

### Behavioral results

There was considerable spread in reading behavior as reported by participants (Fig. 2A). Participants indicated generally liking reading fiction (7-point scale; mean=5.1; median=6; s.d.=1.46). They indicated reading fiction on average about one time per week (5-point scale, 3 = ‘one time per week’; mean=2.7; median=2, s.d.=0.98). The median number of novels read last year was 3 (s.d.=11.6, note the extreme value in Figure 2A which we did not remove since it is a possible value). Participants scored well above chance on the ART (possible scores -12 to 30; mean=8.4, median=8, s.d.=4.4). Scores on the EQ questionnaire were mostly within the normal range (mean=43.9; median=44, s.d.=9.8).

**Figure 2:**
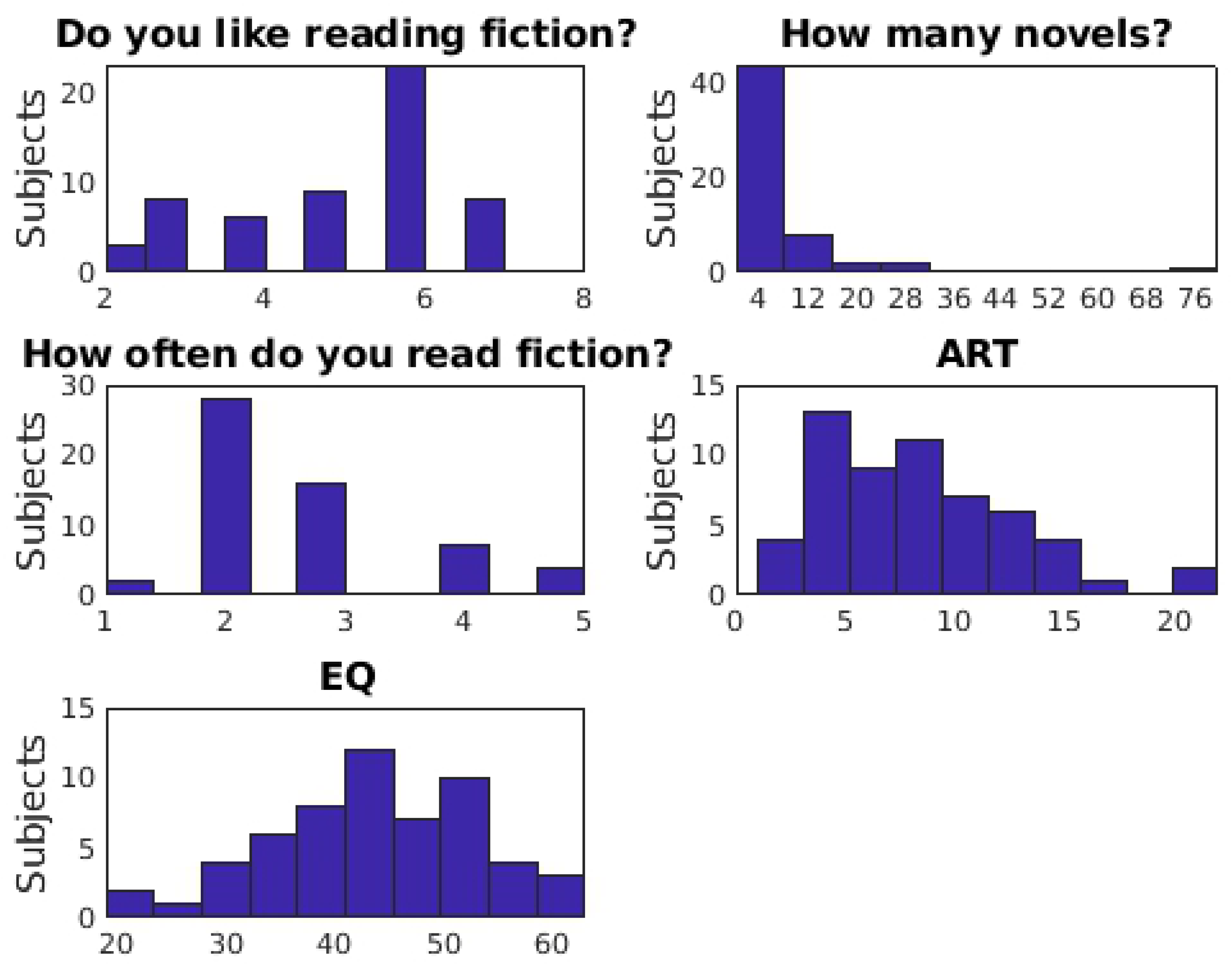
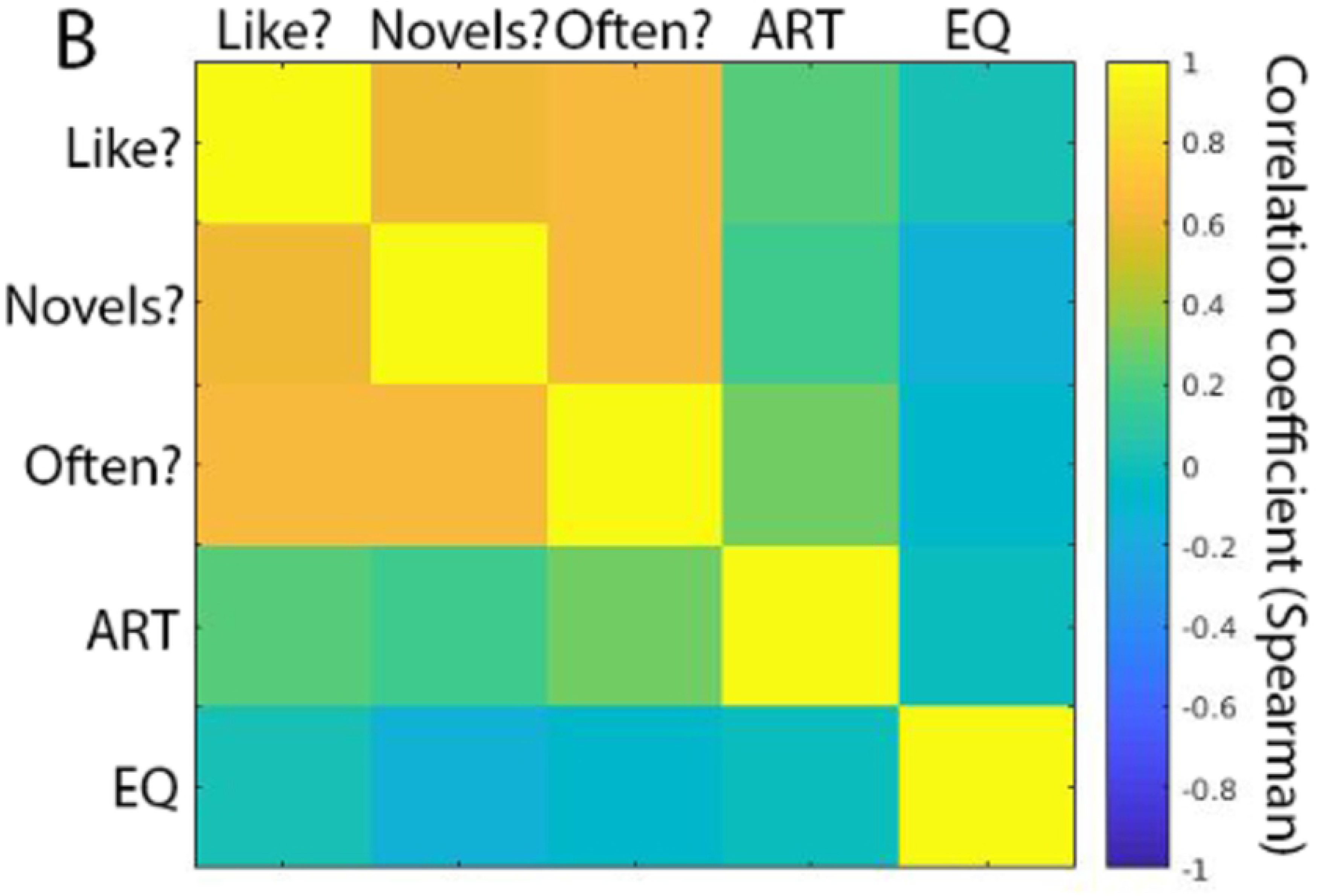
Behavioral Results. A) Histograms of responses to questionnaires. The figure shows the scores on the four measures of reading habits, ART, as well as on the EQ. The plots show that there was considerable spread in all measures within our sample. The extreme value to the How many novels did you read last year (upper right panel) was retained since it is a possible value. B) Spearman correlations between individual difference measures. The three self-report questions concerning reading habits correlated highly with each other (‘Do you like reading fiction?’, ‘How many novels did you read last year?’, and ‘How often do you read fiction?’. The scores on the implicit test of print exposure (Author Recognition Test, ART) were less strongly correlated with the explicit measures, but all correlation values were still sizeable. Correlations with the EQ were low. See text for correlation values. Note that all correlation coefficients are Spearman’s Rank correlations that are robust against outliers.

The reading habits questions correlated highly with each other. Pairwise Spearman correlations between the questions ‘Do you like reading fiction?’, ‘How often do you read fiction?’, and ‘How many novels did you read last year?’ were all high (all rho(56)>0.62, all p<0.001; Fig. 2B). The ART correlated moderately with the reading habits self-reports (Fig. 2B; ‘Do you like reading fiction?’:Rho=0.22, p=0.09; ‘How often do you read fiction?’:Rho=0.16, p=0.22; ‘How many novels did you read last year?’:Rho=0.30, p=0.02). The size of the correlation coefficient between ART and self-reported amount of reading is comparable with that observed in a much larger sample by Acheson and colleagues (Acheson et al., 2008). The ART was mainly developed as an objective measure of print exposure in the sense that it is not influenced by response bias. We are, however, not concerned that the self-report measures (‘How often do you read fiction?’, etc.) suffered from social desirability in the current sample, since participants did not feel inhibited to report that they hardly ever read (see Fig. 2A). While the reading habits questions target more current lifestyle habits, the ART questionnaire captures lifetime exposure to fiction better, but does not necessarily reflect current habits. Given that the ART also has its shortcomings (one can score very high on the ART without actually reading books), we had no reason to value the ART higher above the self-report measures, or vice versa. The EQ questionnaire did not correlate significantly with any of the other measures (all rho<|0.13|, all p>0.3).

Most important for the present individual differences approach is that there was considerable spread in all measures, and most notably in those measuring amount of fiction reading.

#### fMRI network results

The control analysis examined the connection strengths depending on trait empathy. It showed a set of areas with statistically higher node degree of correlations between regions and the scores of participants on the ***EQ*** questionnaire. These included (among others) several regions in inferior temporal cortex including fusiform cortex, the anterior medial prefrontal cortex (MPFC) and orbitofrontal cortex, temporo-parietal junction, and the right frontal pole (see Table 2, Figure 3).

**Figure 3:**
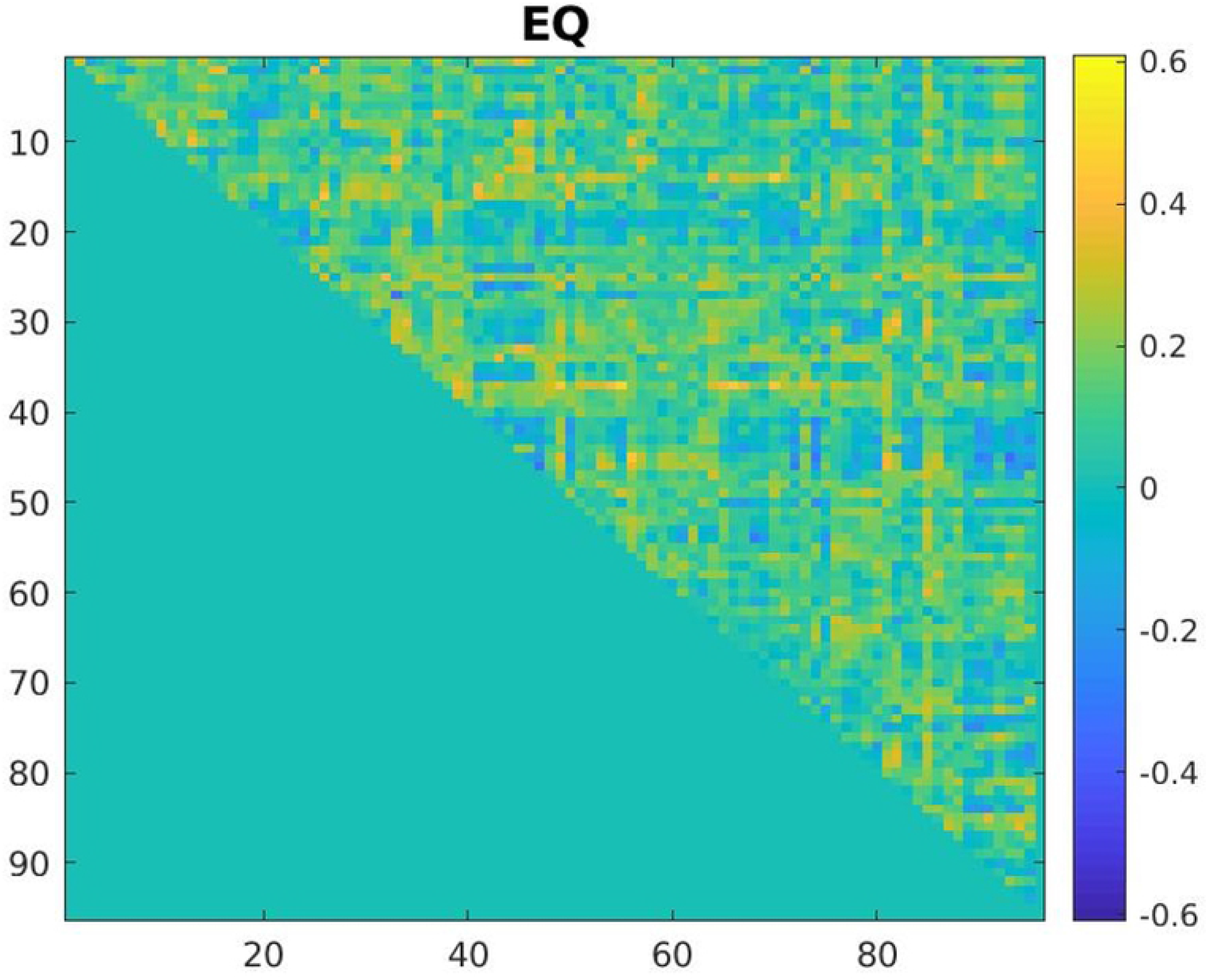
Correlation matrix of the region to region connections from all 96 cortical areas in the Harvard-Oxford atlas, covering the whole cortical sheet (for legend see Supplementary Materials S3) modulated by the EQ score.

**Table 2:** Empathy Quotient (EQ)

| Region | Connections with: |
| --- | --- |
| L anterior Inferior Temporal Gyrus (8 connections) | L Parietal Operculum Cortex |
|  | R Frontal Pole |
|  | R Middle Frontal Gyrus |
|  | R Inferior Frontal Gyrus, opercularis |
|  | R Inferior Temporal Gyrus, temporooccipital |
|  | R Superior Parietal Lobule |
|  | R anterior Supramarginal Gyrus |
|  | R superior lateral occipital cortex |
| L Inferior Temporal Gyrus, temporooccipital (9 connections) | L Supplementary Motor Area |
|  | L Cuneal Cortex |
|  | L Frontal Operculum Cortex |
|  | L Central Opercular Cortex |
|  | R Insula |
|  | R anterior Cingulate Gyrus |
|  | R Cuneal Cortex |
|  | R Central Opercular Cortex |
|  | L Insula |
| L Frontal Medial Cortex (12 connections) | L Supplementary Motor Area |
|  | L Cuneal Cortex |
|  | L Central Opercular Cortex |
|  | R Insula |
|  | R Supplementary Motor Area |
|  | R Cuneal Cortex |
|  | R posterior Parahippocampal Gyrus |
|  | R Occipital Fusiform Gyrus |
|  | R Planum Polare |
|  | L Insula |
|  | L Precentral Gyrus |
|  | L Postcentral Gyrus |
| L anterior Temporal Fusiform Gyrus (16 connections) | L Temporal Occipital Fusiform Cortex |
|  | L Occipital Pole |
|  | R Frontal Pole |
|  | R Superior Frontal Gyrus |
|  | R Middle Frontal Gyrus |
|  | R Inferior Frontal Gyrus, triangularis |
|  | R Inferior Frontal Gyrus, opercularis |
|  | R Precentral Gyrus |
|  | R Inferior Temporal Gyrus, temporooccipital |
|  | R Postcentral Gyrus |
|  | R Superior Parietal Lobule |
|  | R anterior Supramarginal Gyrus |
|  | R superior lateral occipital cortex |
|  | R Frontal Medial Cortex |
|  | R Supplementary Motor Area |
|  | L Precentral Gyrus |
| L Heschl's Gyrus's Gyrus (8 connections) | R Frontal Pole |
|  | R Temporal Pole |
|  | R Frontal Orbital Cortex |
|  | L Frontal Pole |
|  | L Temporal Pole |
|  | L anterior Middle Temporal Gyrus |
|  | L Middle Temporal Gyrus, temporooccipital |
|  | L Frontal Orbital Cortex |
| L Planum Temporale (10 connections) | R Inferior Frontal Gyrus, triangularis |
|  | R Temporal Pole |
|  | R Frontal Orbital Cortex |
|  | L Frontal Pole |
|  | L Temporal Pole |
|  | L anterior Superior Temporal Gyrus |
|  | L anterior Middle Temporal Gyrus |
|  | L posterior Middle Temporal Gyrus |
|  | L Middle Temporal Gyrus, temporooccipital |
|  | L Frontal Orbital Cortex |
| R Frontal Pole (9 connections) | R Precentral Gyrus |
|  | R Precuneus |
|  | L Insula |
|  | L anterior Inferior Temporal Gyrus |
|  | L Supplementary Motor Area |
|  | L Precuneus |
|  | L anterior Parahippocampal Gyrus |
|  | L anterior Temporal Fusiform Cortex |
|  | L Heschl's Gyrus |
| R Temporal Fusiform, ant (12 connections) | R Parietal Operculum Cortex |
|  | L anterior Superior Temporal Gyrus |
|  | L posterior Middle Temporal Gyrus |
|  | L Angular Gyrus |
|  | L posterior Cingulate Gyrus |
|  | L Precuneus |
|  | L Supracalcarine Cortex |
|  | R Superior Frontal Gyrus |
|  | R Inferior Frontal Gyrus, triangularis |
|  | R posterior Middle Temporal Gyrus |
|  | R Frontal Medial Cortex |
|  | R Paracingulate Gyrus |

There was a statistically significant difference in node degree in a set of regions depending on how participants scored on the question ‘***How often do you read fiction****?*’ (Table 3, Fig. 4, Figure 6A). In these regions, the number of region-region by individual difference correlations was statistically higher than expected by chance. This effect was present in a set of regions including (among others) bilateral inferior frontal sulci, bilateral anterior middle temporal regions, anterior medial prefrontal cortex (MPFC),Right posterior supramarginal gyrus, the lingual gyri bilaterally, and the middle frontal cortex bilaterally (see Table 3, Fig 4, Figure 6A). The effect was also present in several regions in sensory and motor cortices.

**Figure 4:**
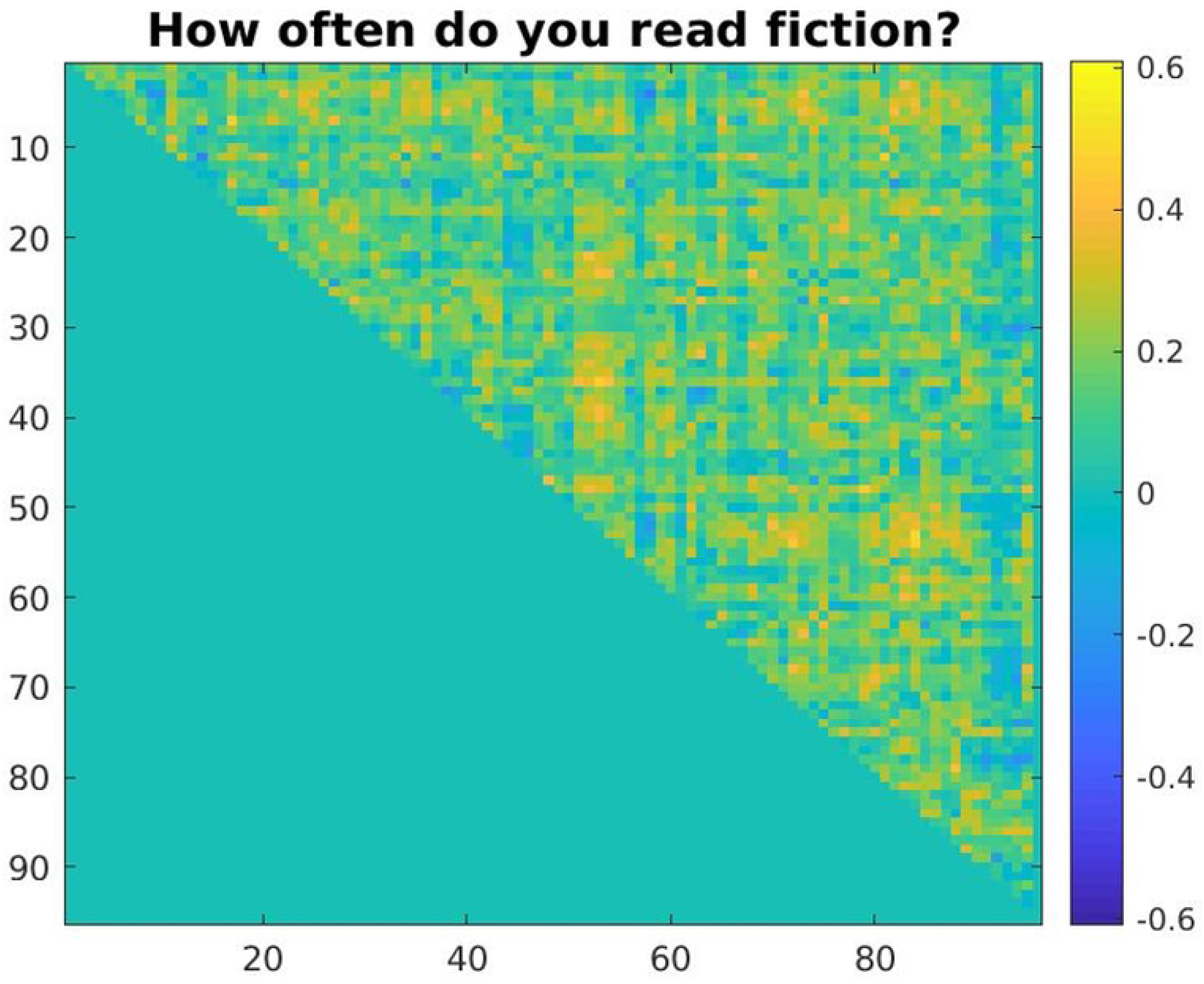
Correlation matrix of the region to region connections from all 96 cortical areas in the Harvard-Oxford atlas, covering the whole cortical sheet (for legend see Supplementary Materials S3) modulated by how high participants scored on the questions ‘How often do you read fiction?’.

**Figure 5:**
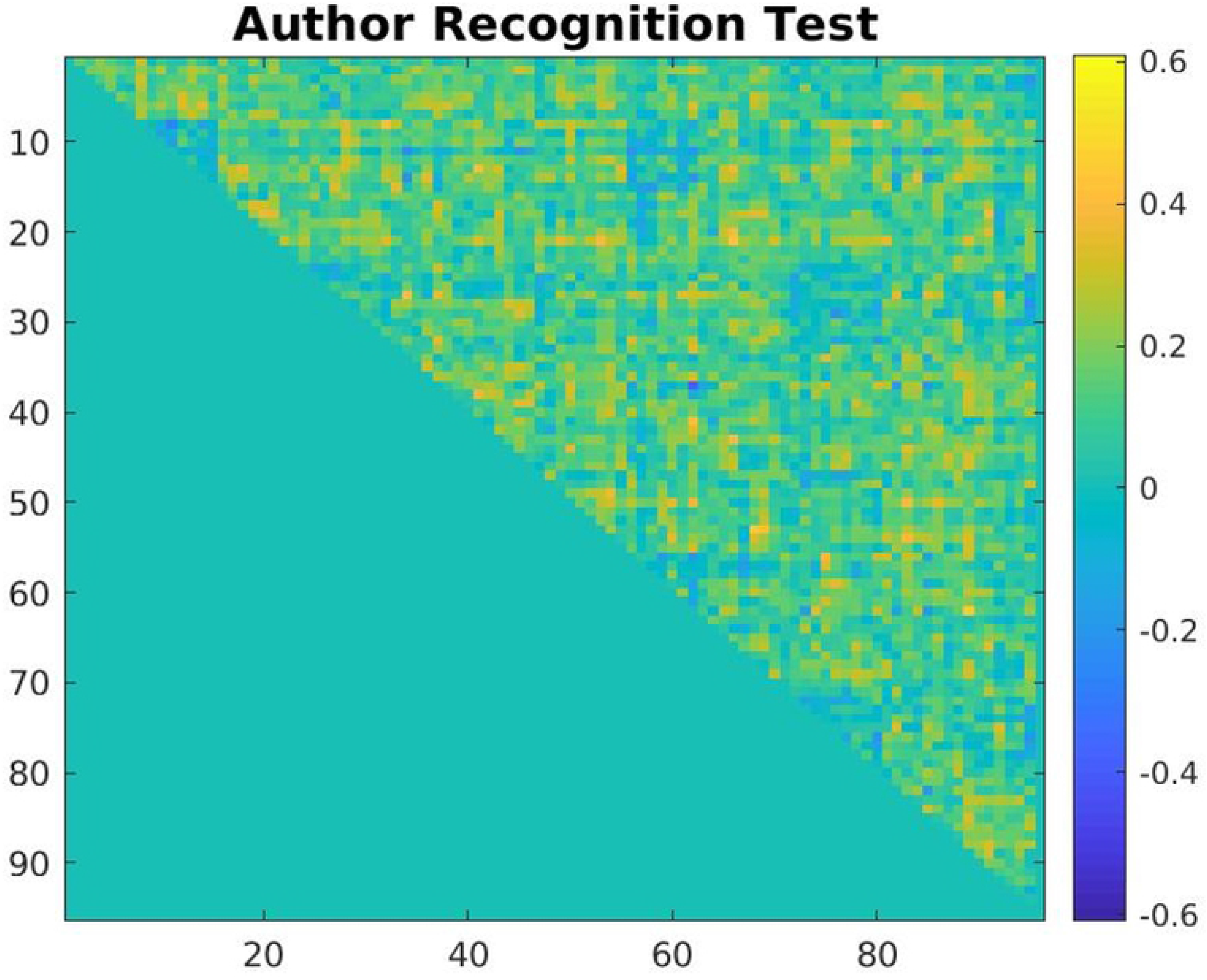
Correlation matrix of the region to region connections from all 96 cortical areas in the Harvard-Oxford atlas, covering the whole cortical sheet (for legend see Supplementary Materials S2) modulated by the ART score.

**Figure 6:**
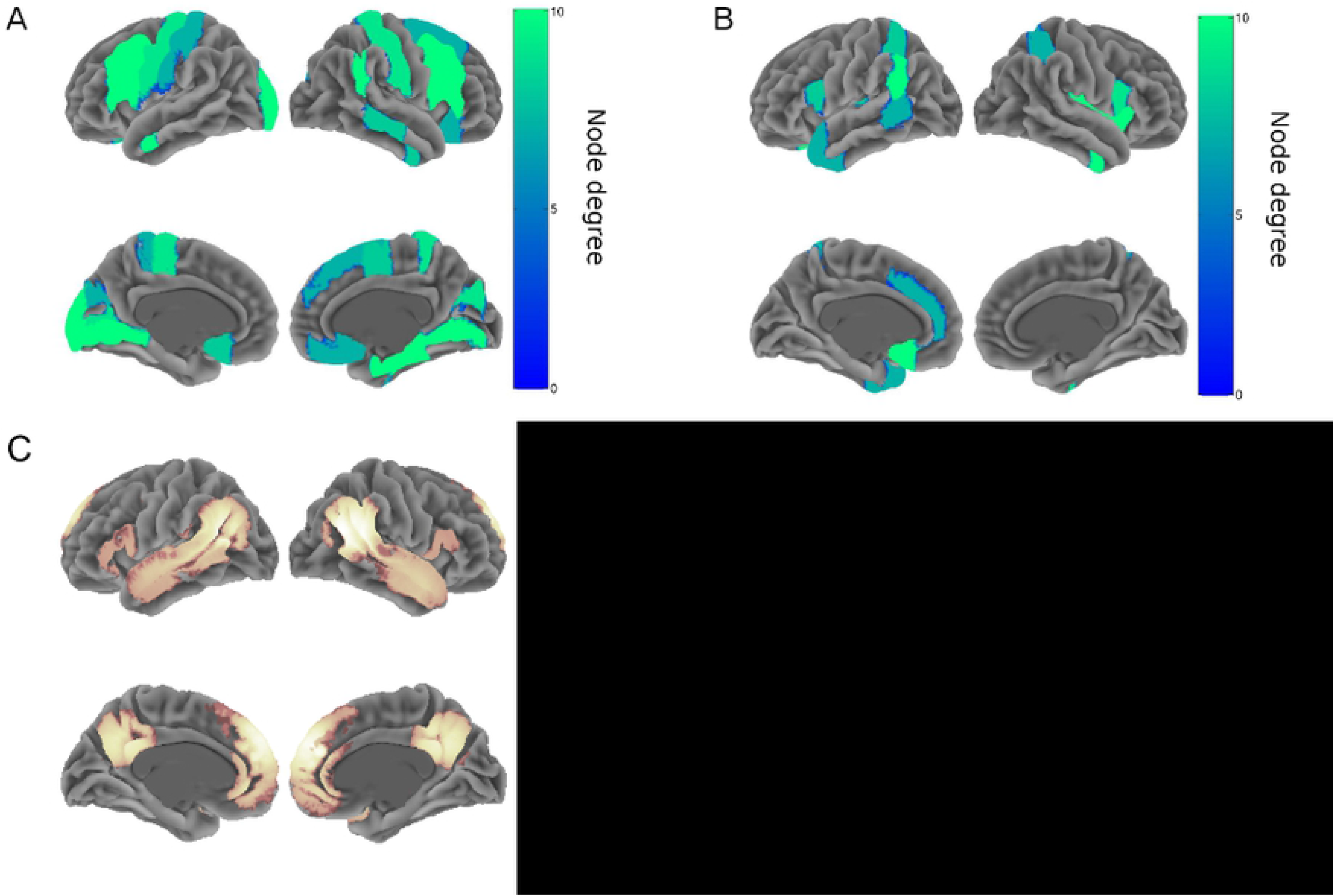
fMRI network results. Displayed are regions which had statistically significantly higher number of connections with other regions (node degree), depending on an individual difference measure. For instance, in A)Regions are displayed in which participants who scored higher on ‘How often do you read fiction?’ had significantly more connections with other regions during listening to the narratives as compared to the reversed speech baseline. B)Results for correlations with Author Recognition Test (ART). C) Shows the BOLD results of a meta-analysis of ToM tasks by Schurz et al 2013 (displayed is a newer more accurate analysis of the data by the author).

**Table 3:**
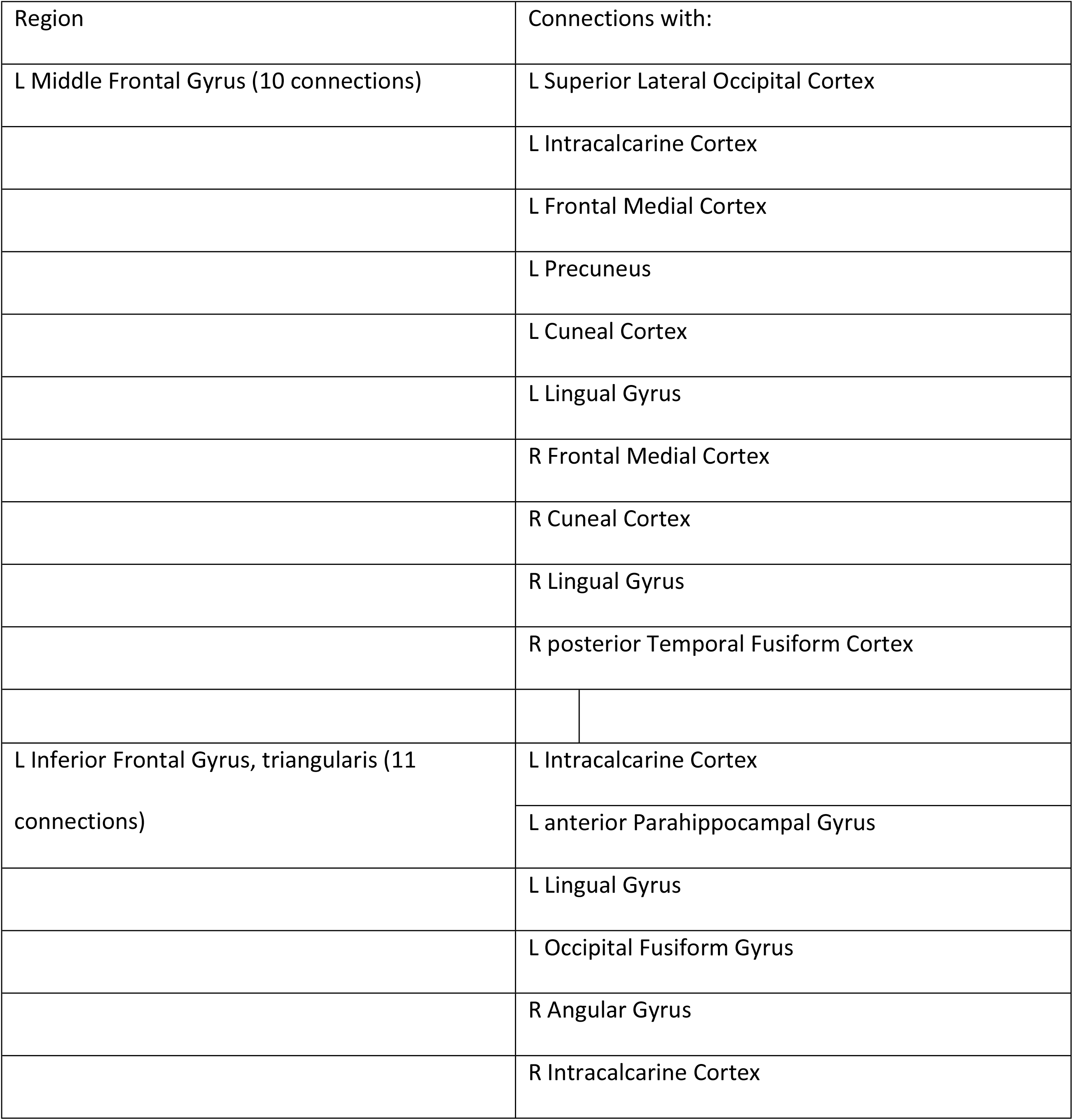
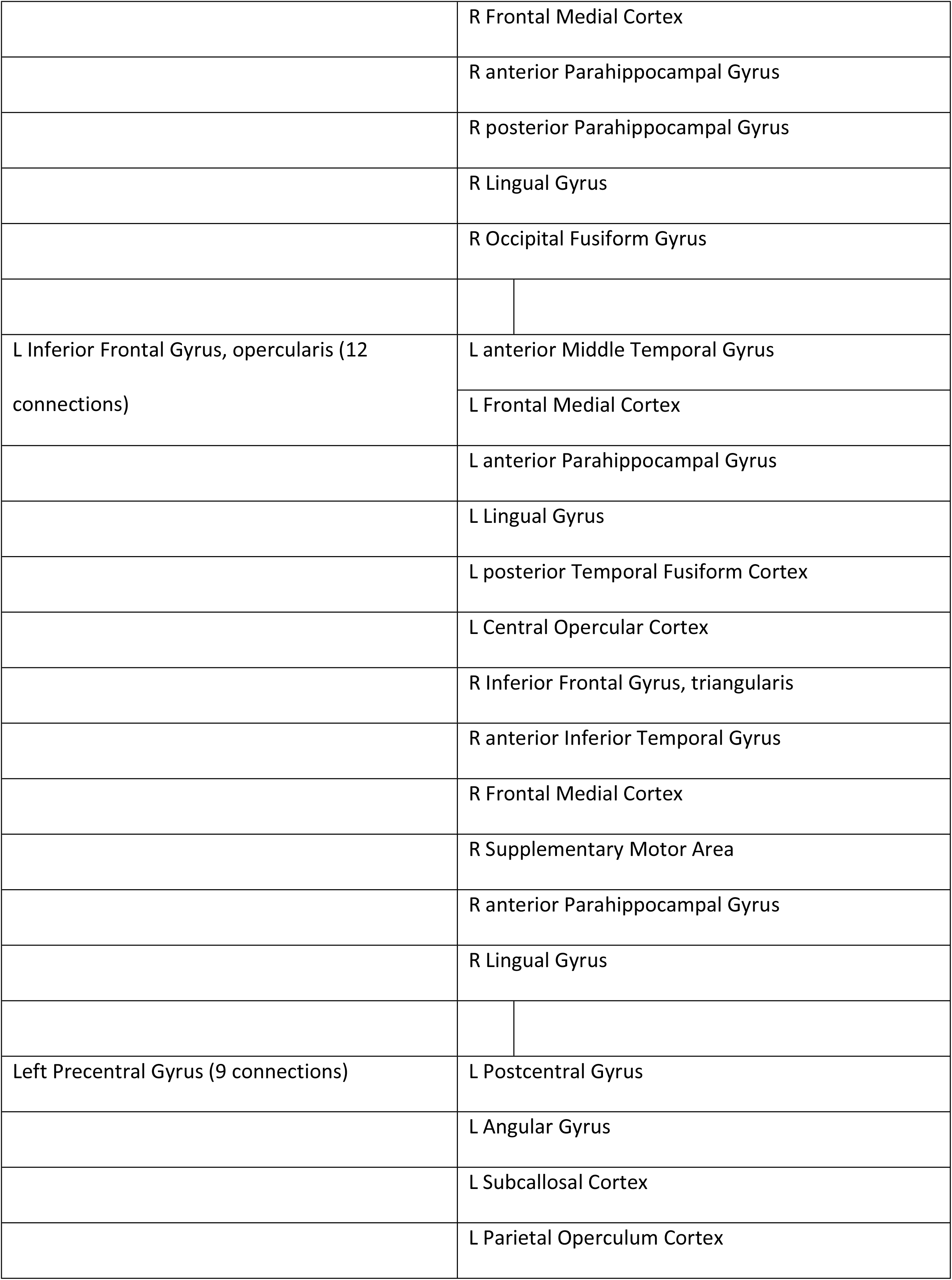
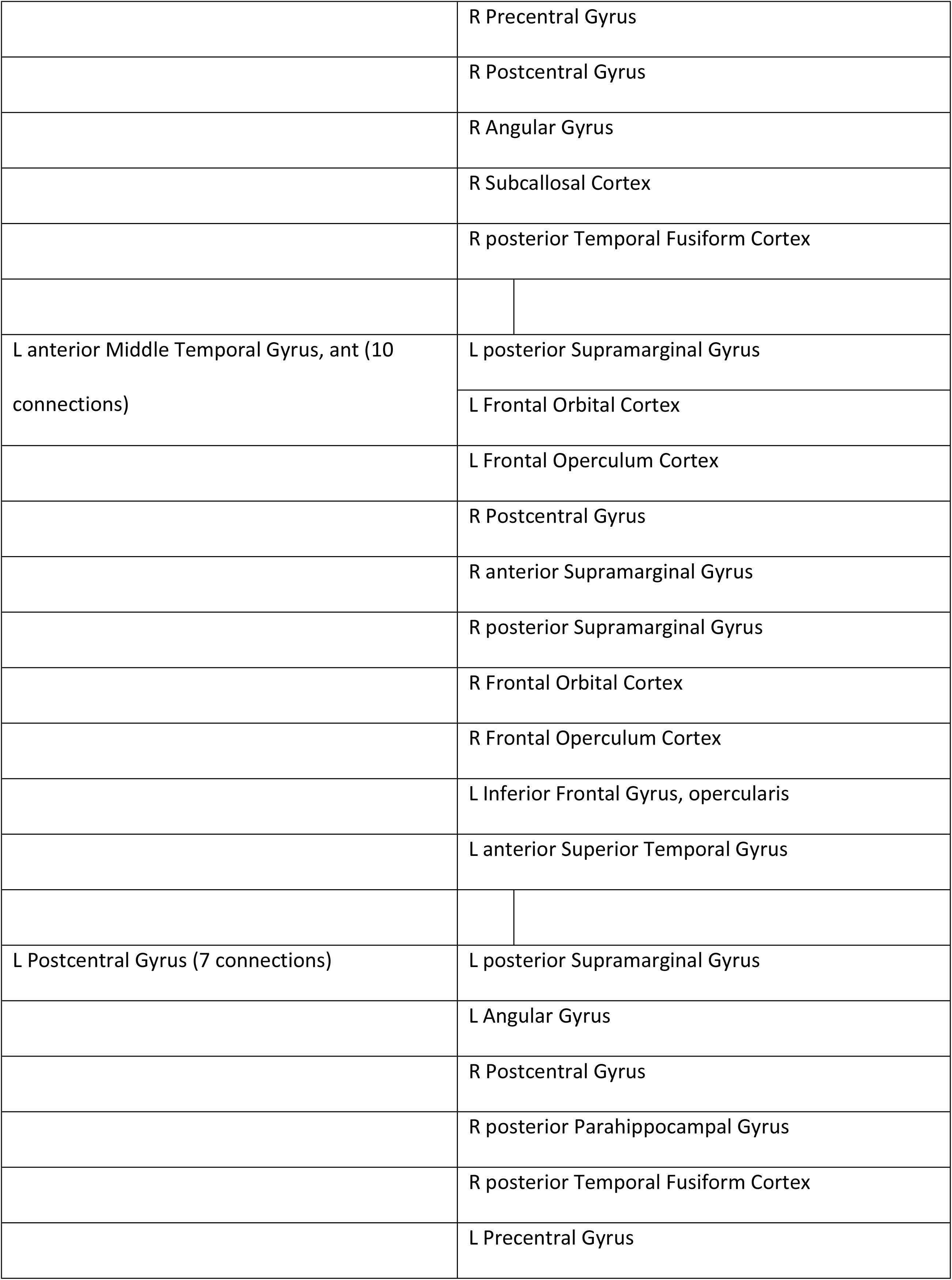
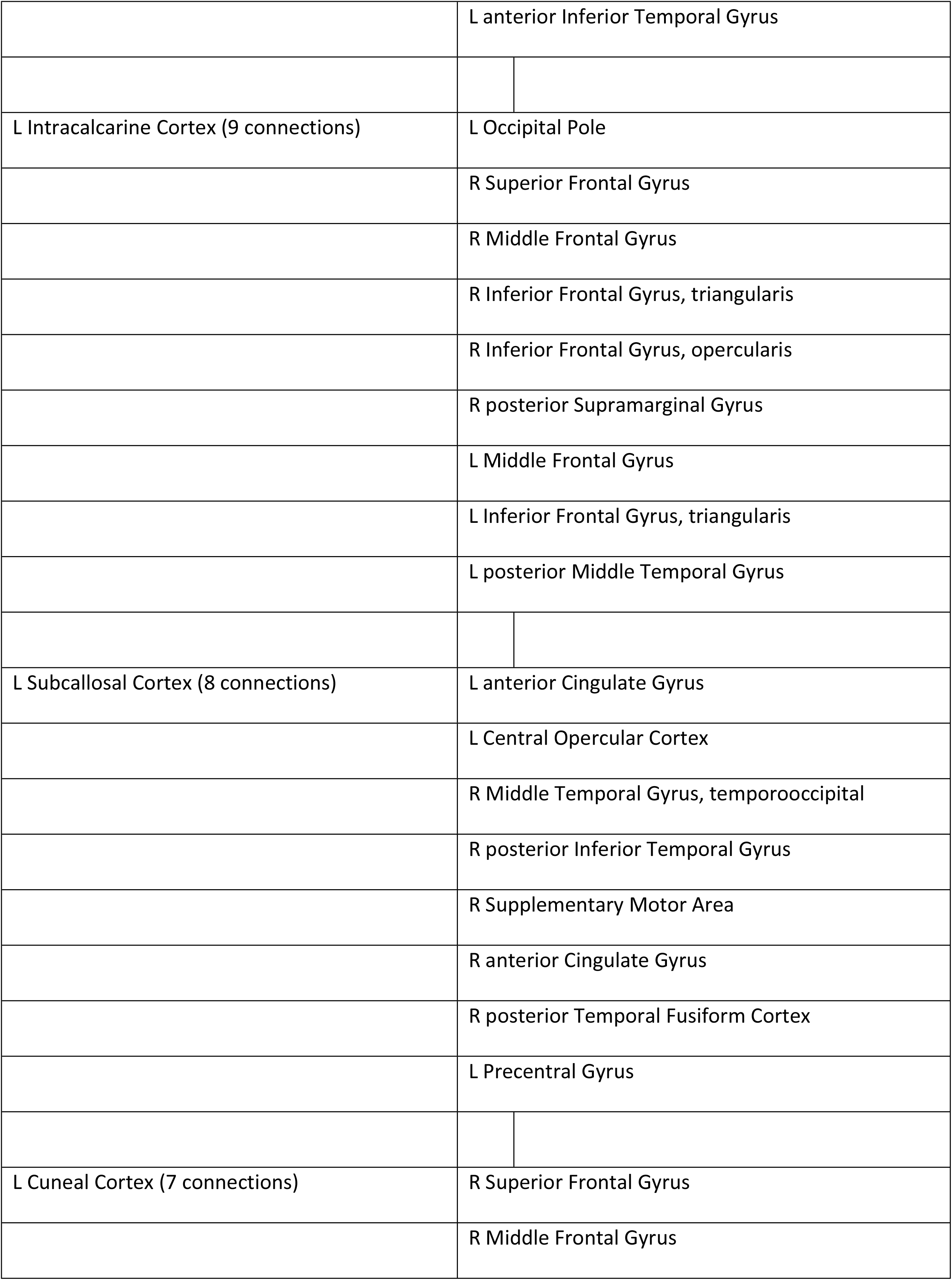
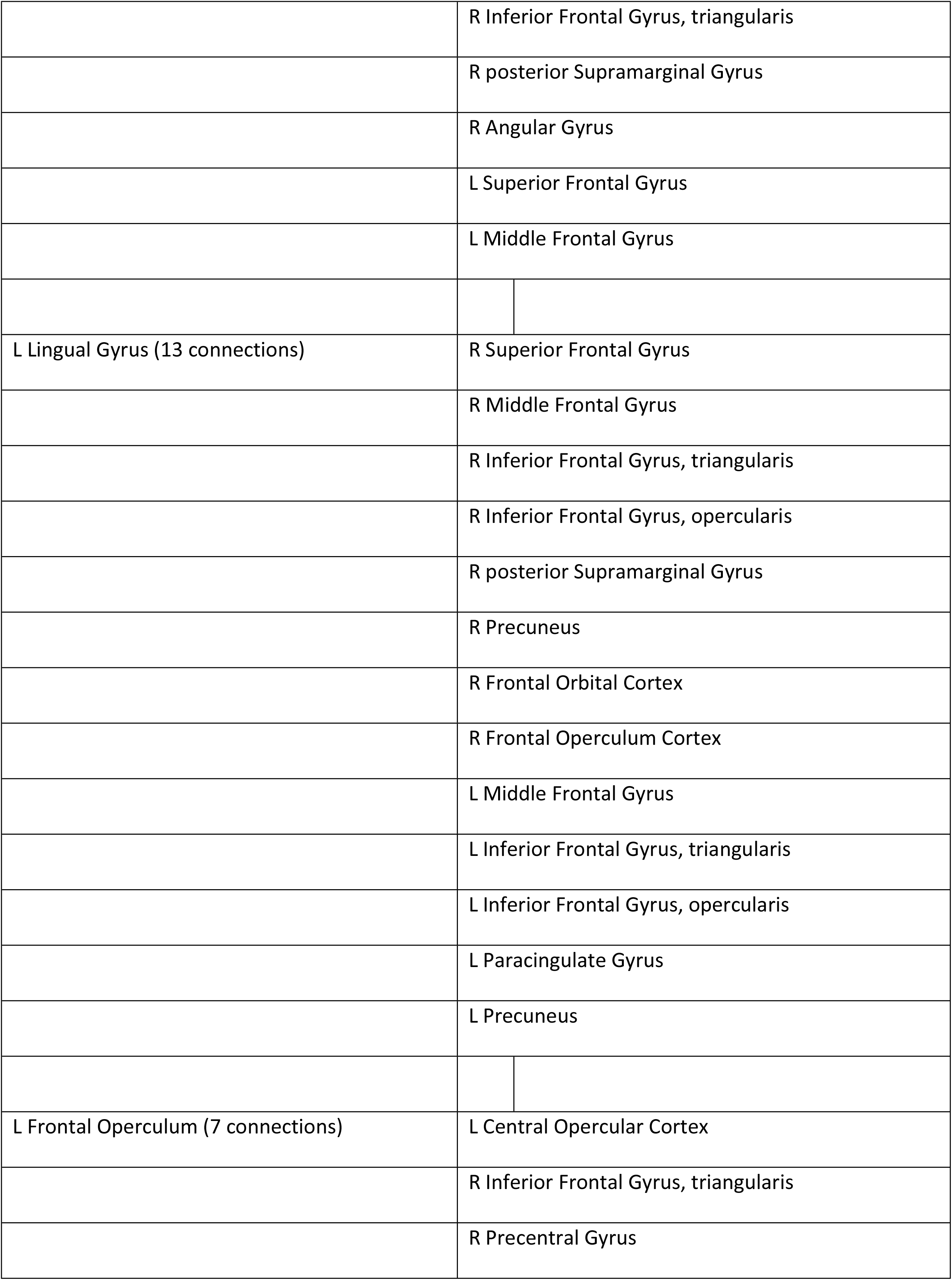
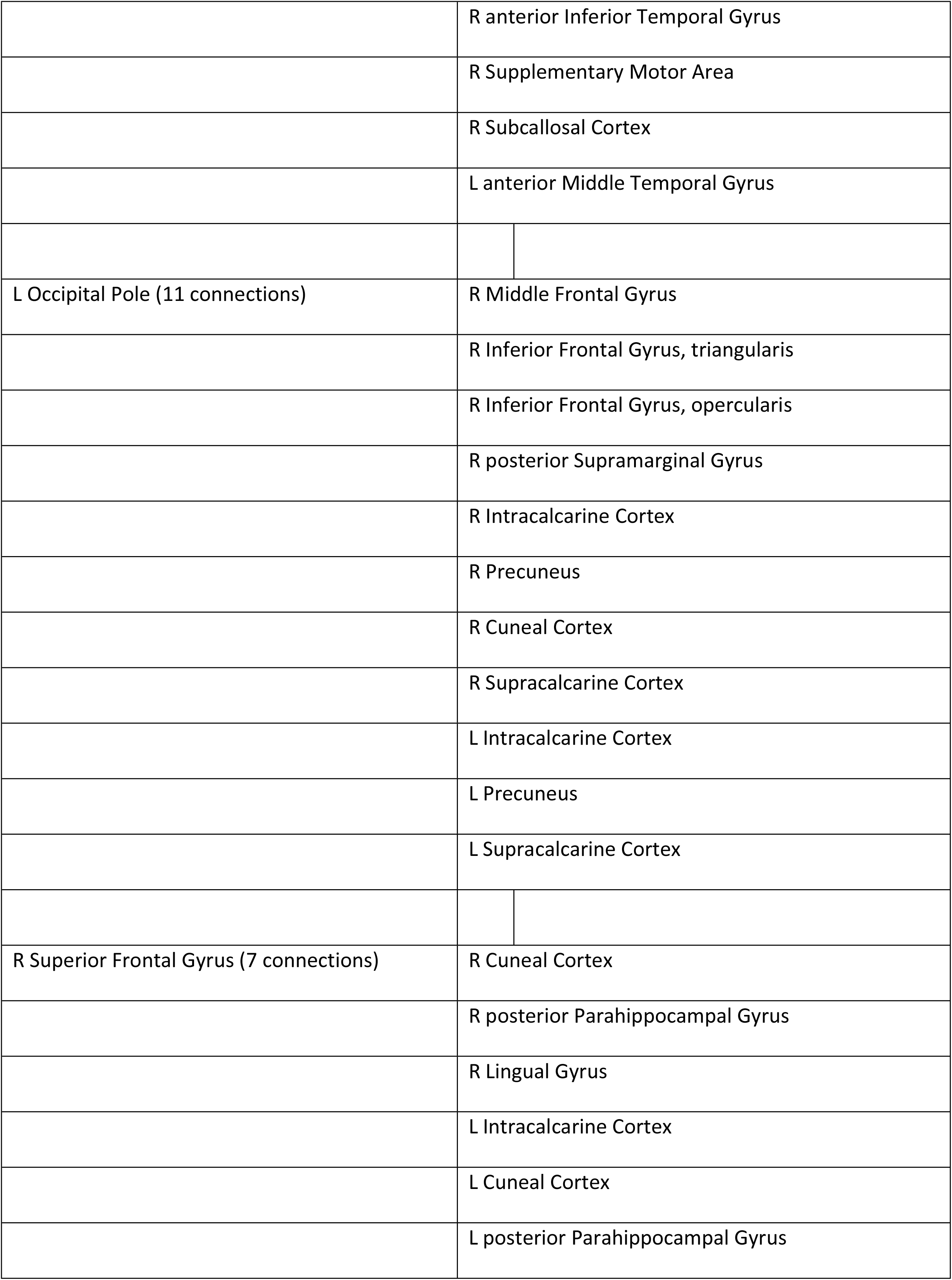
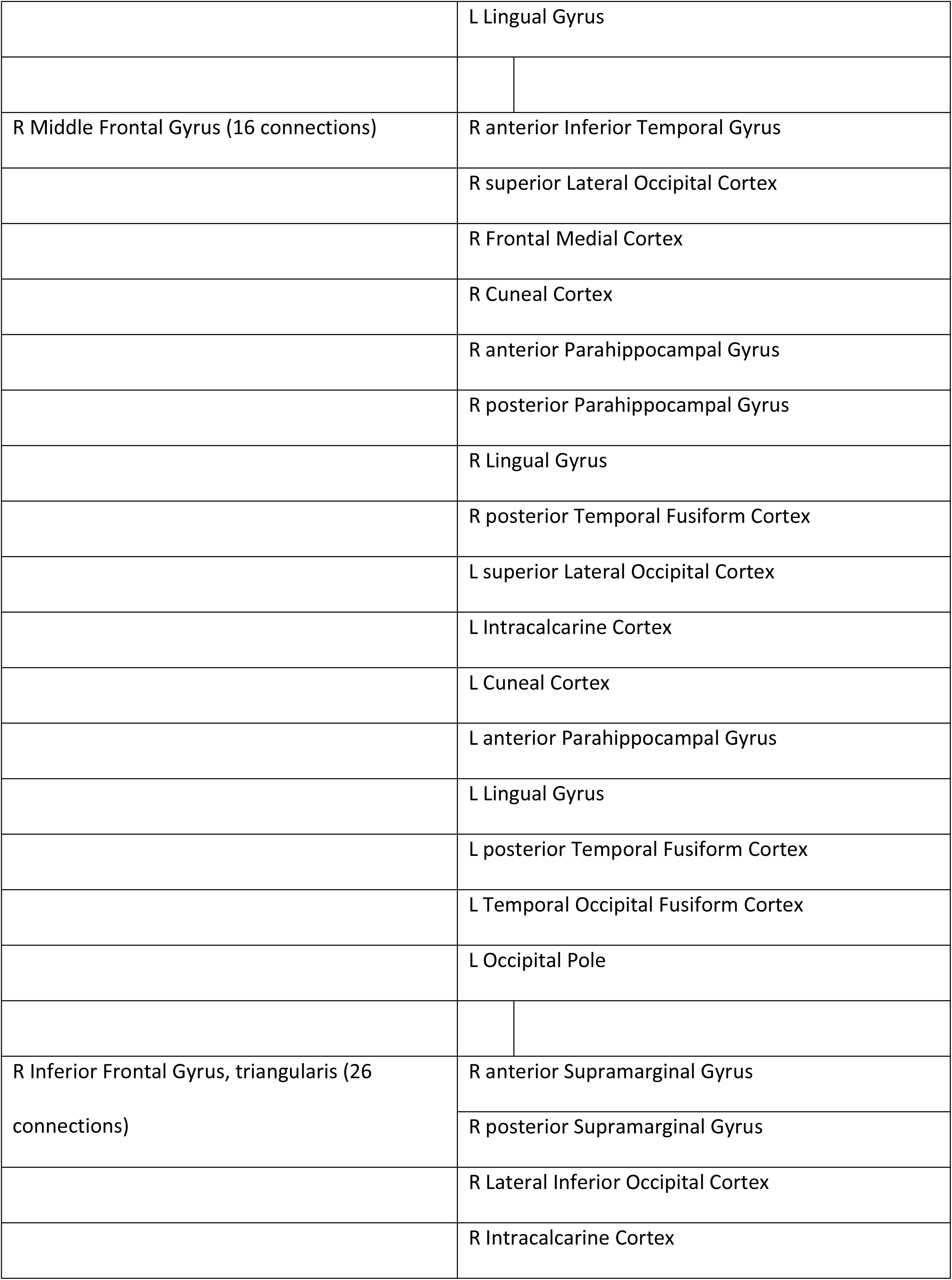
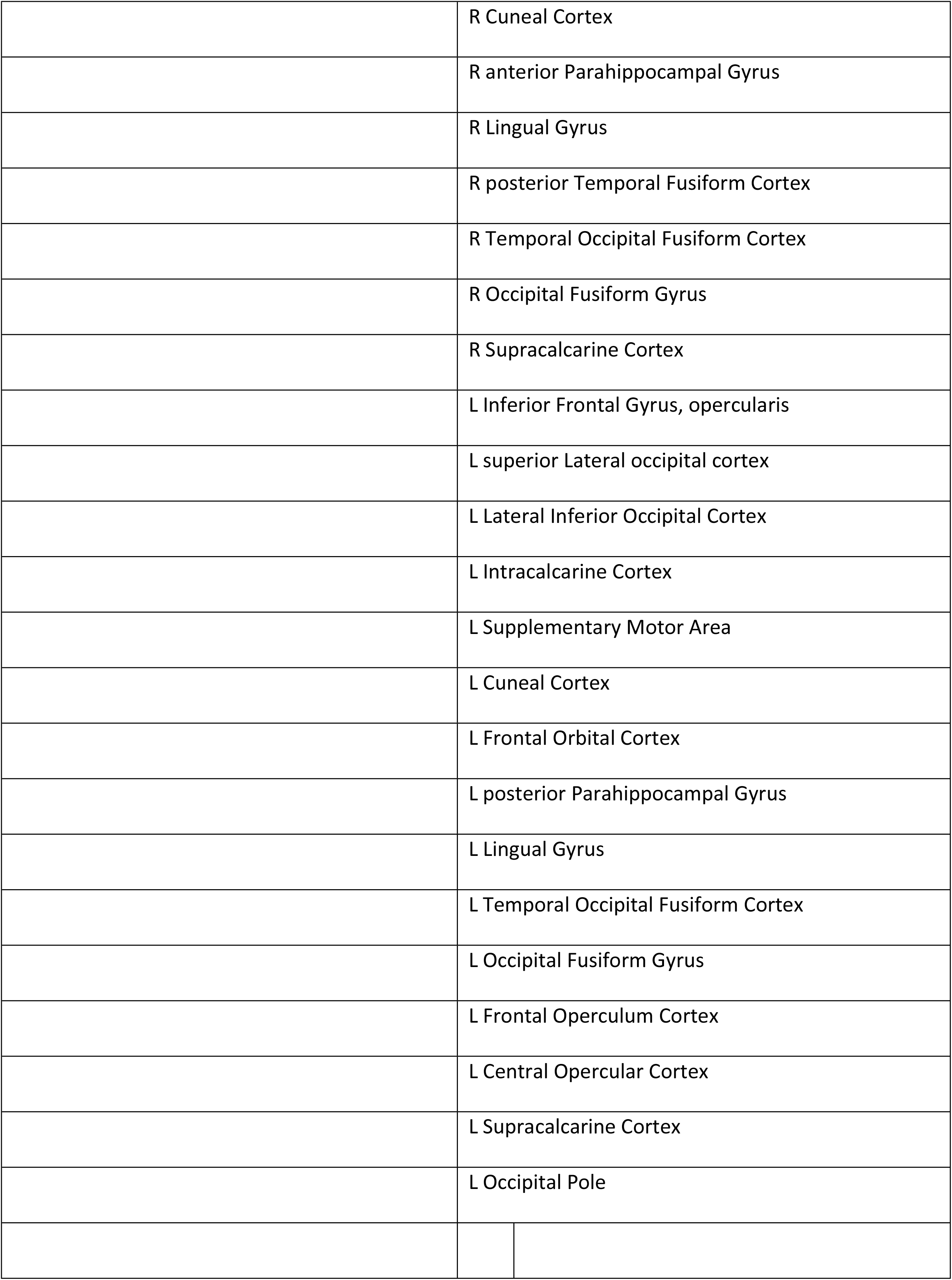
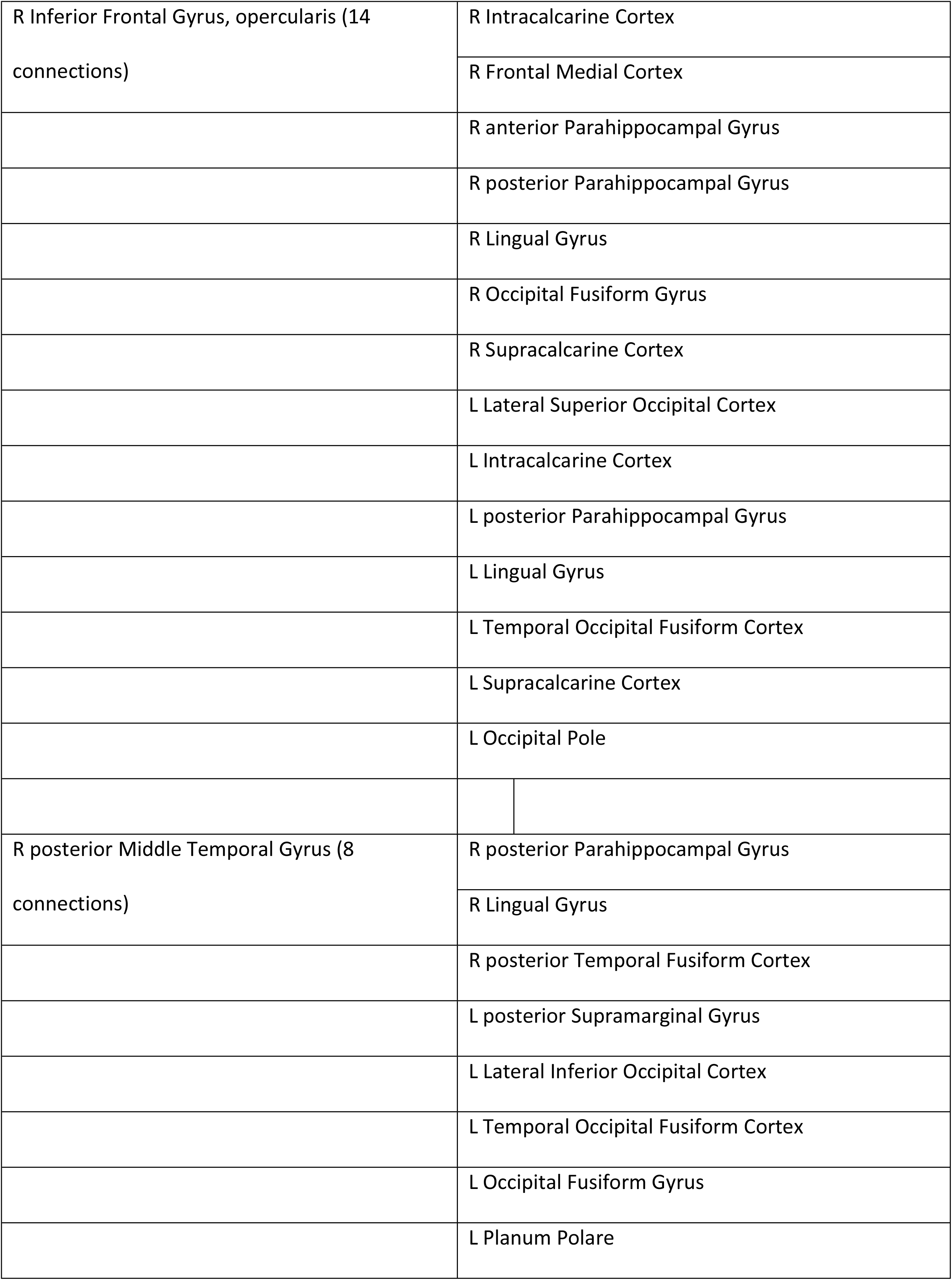
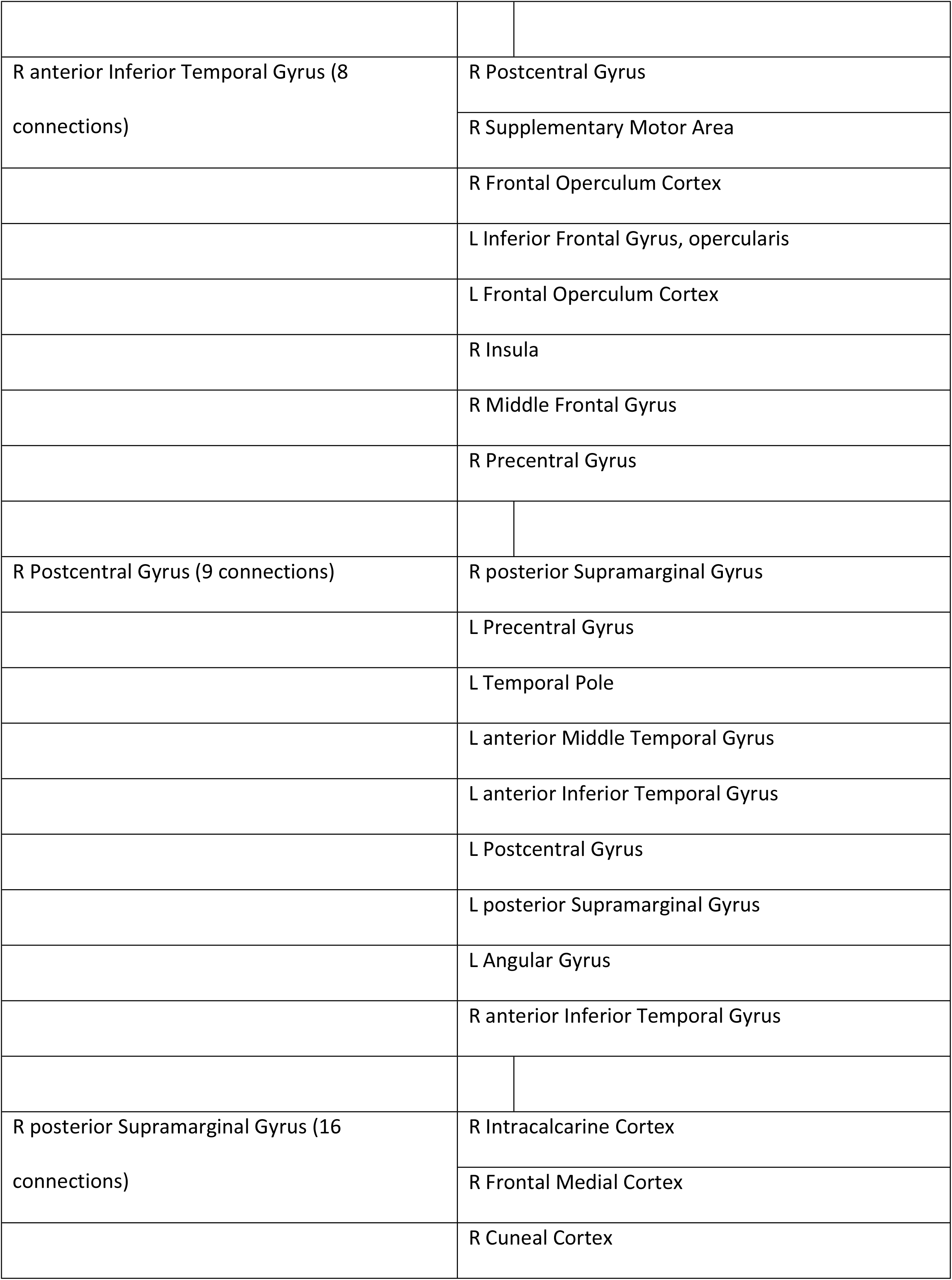
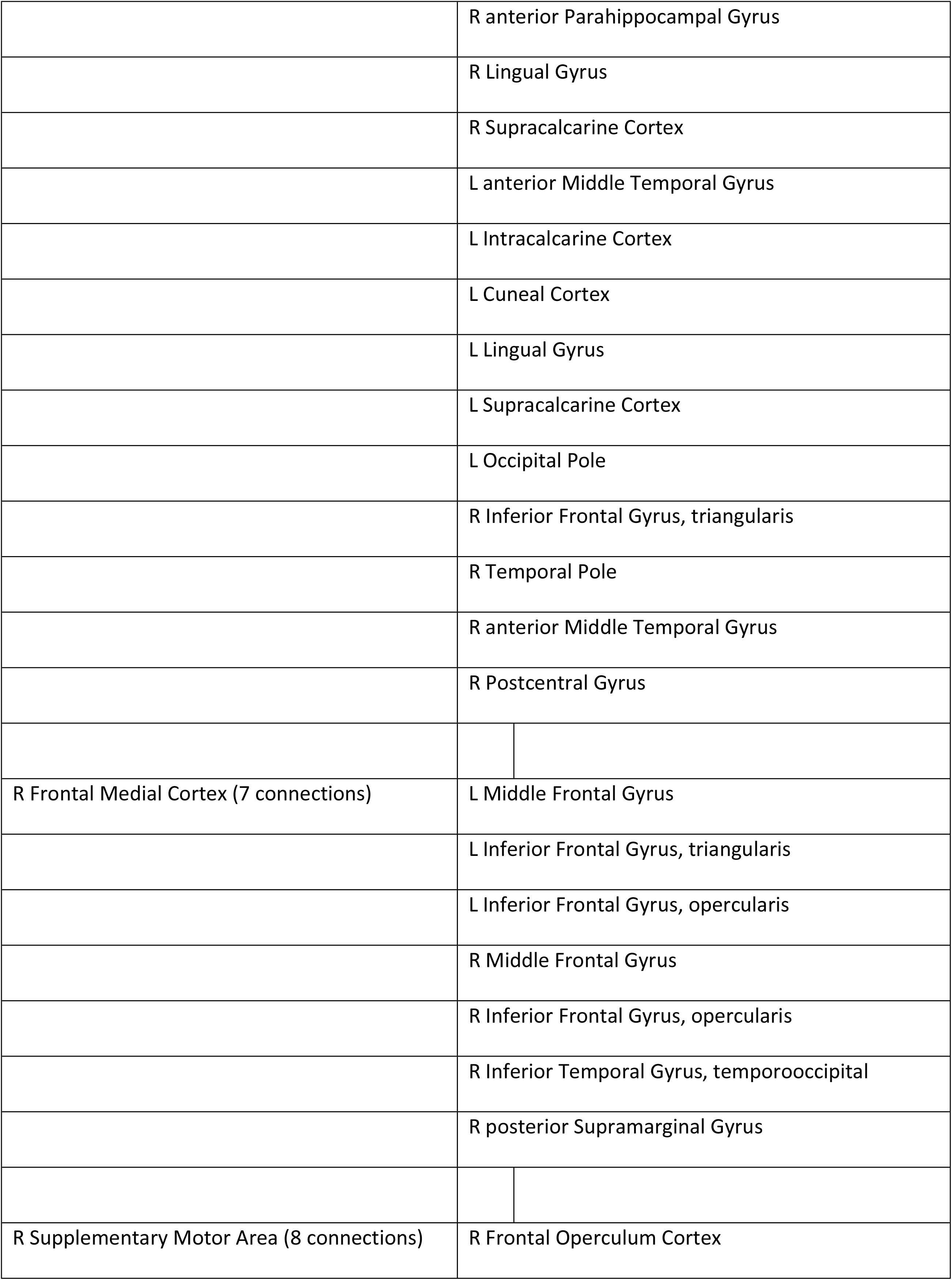
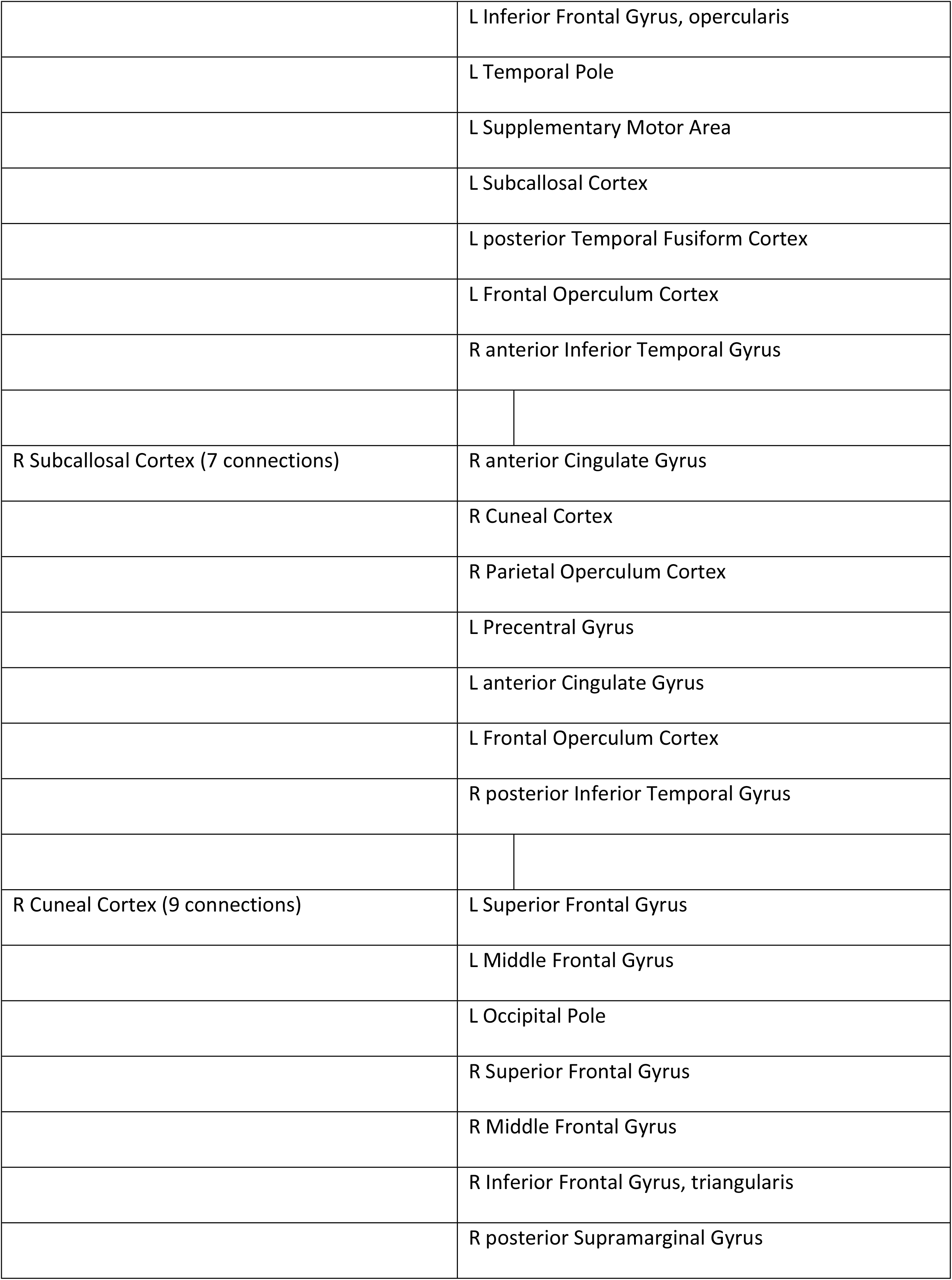
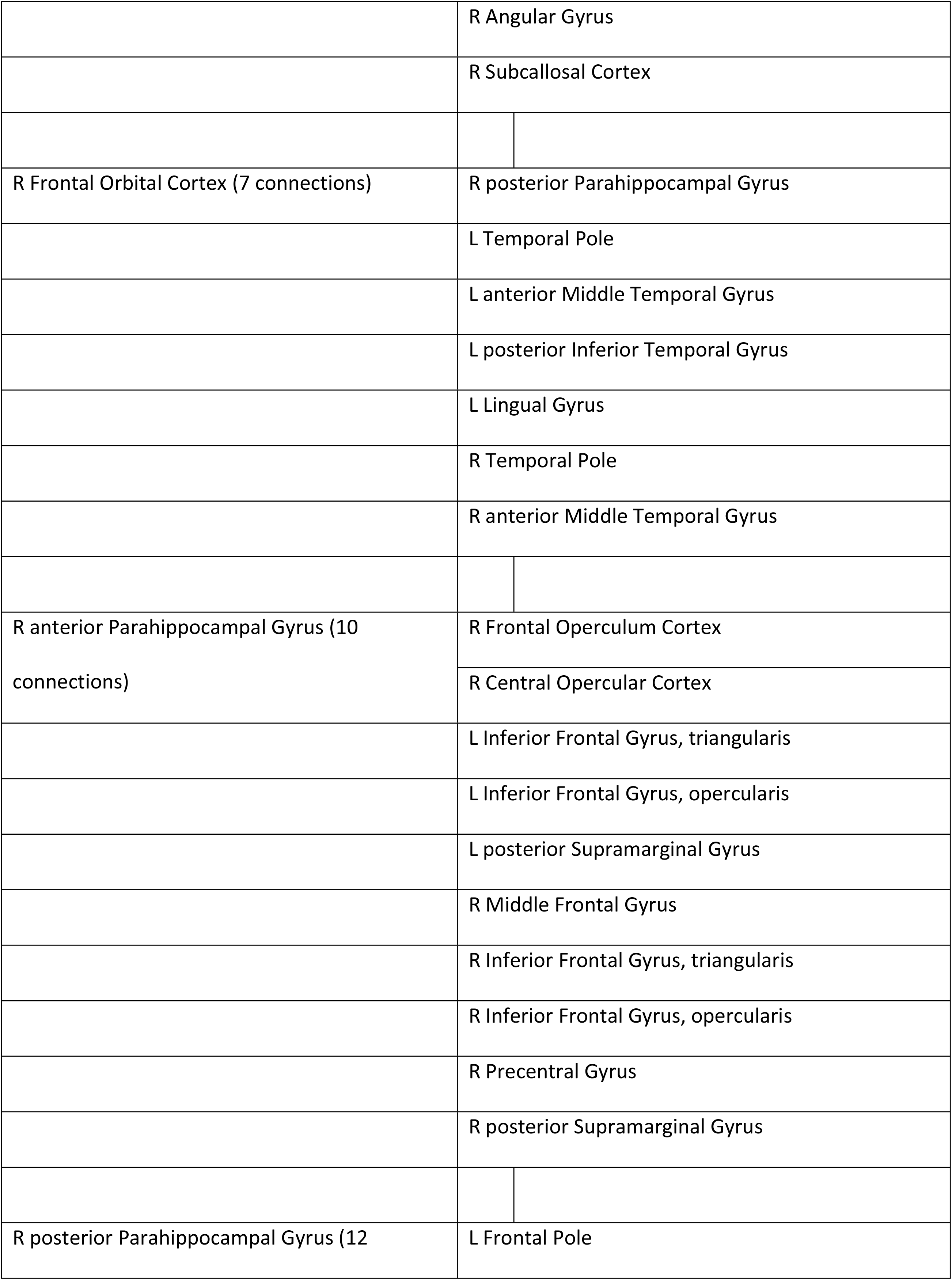
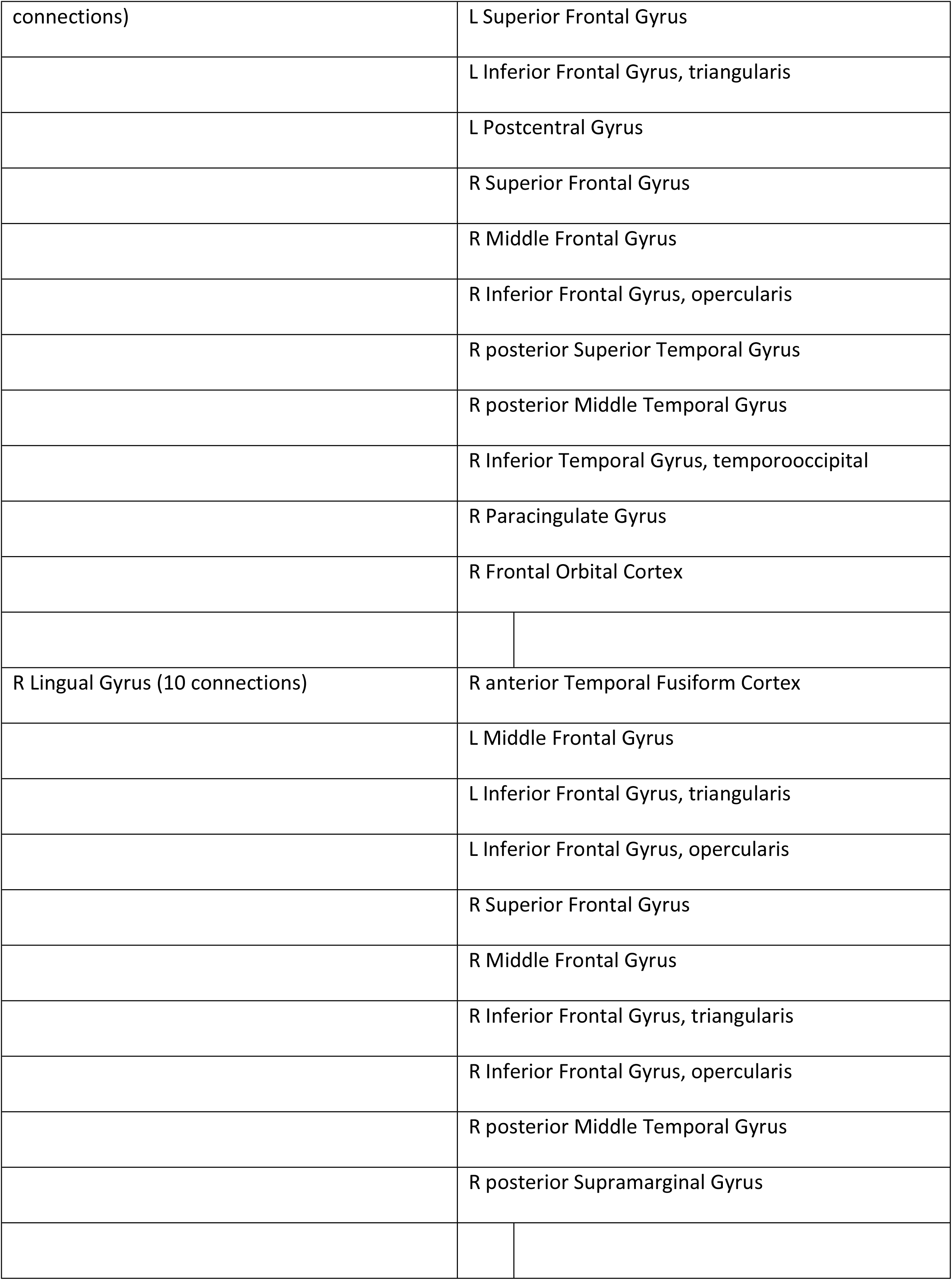
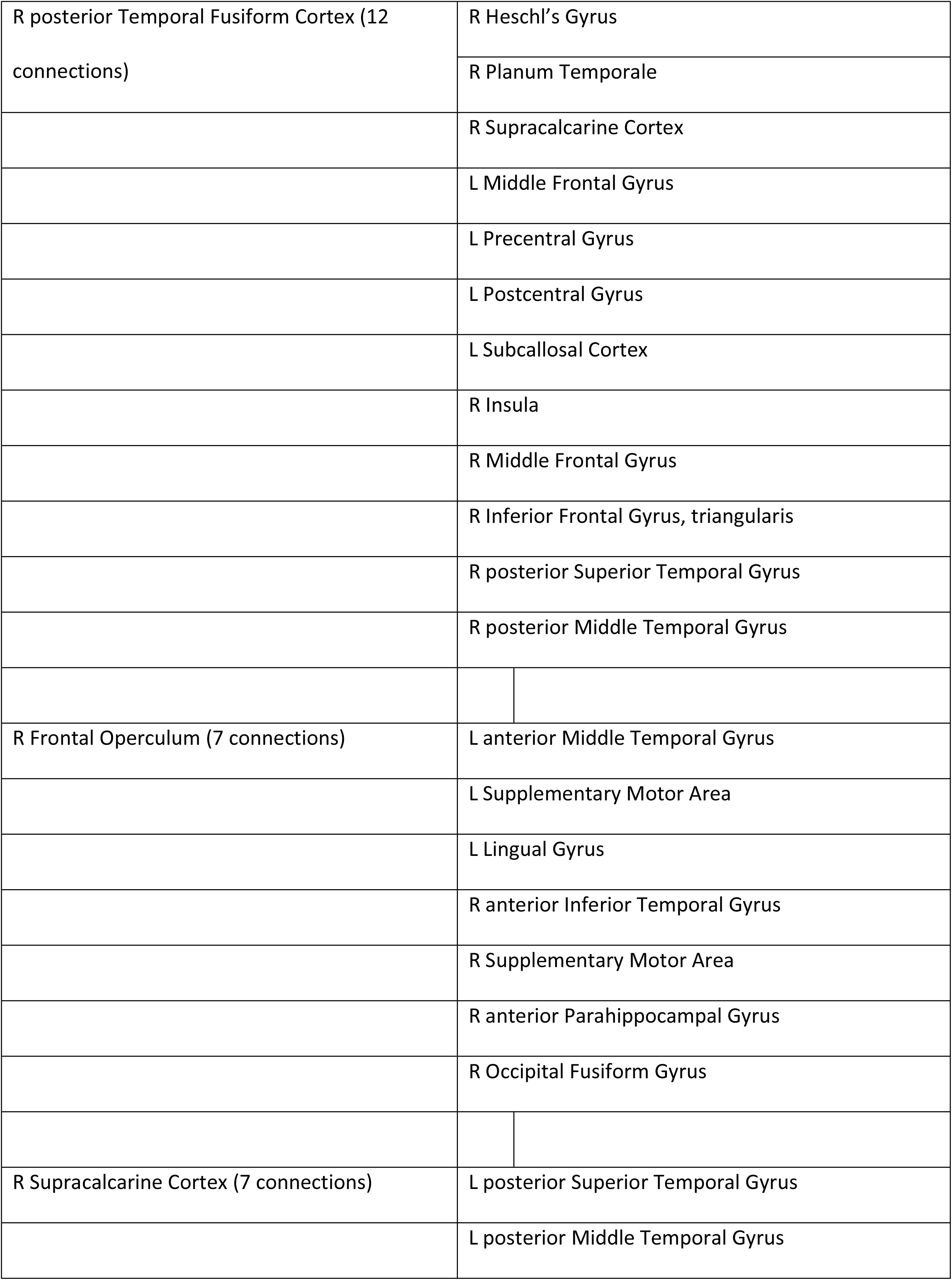
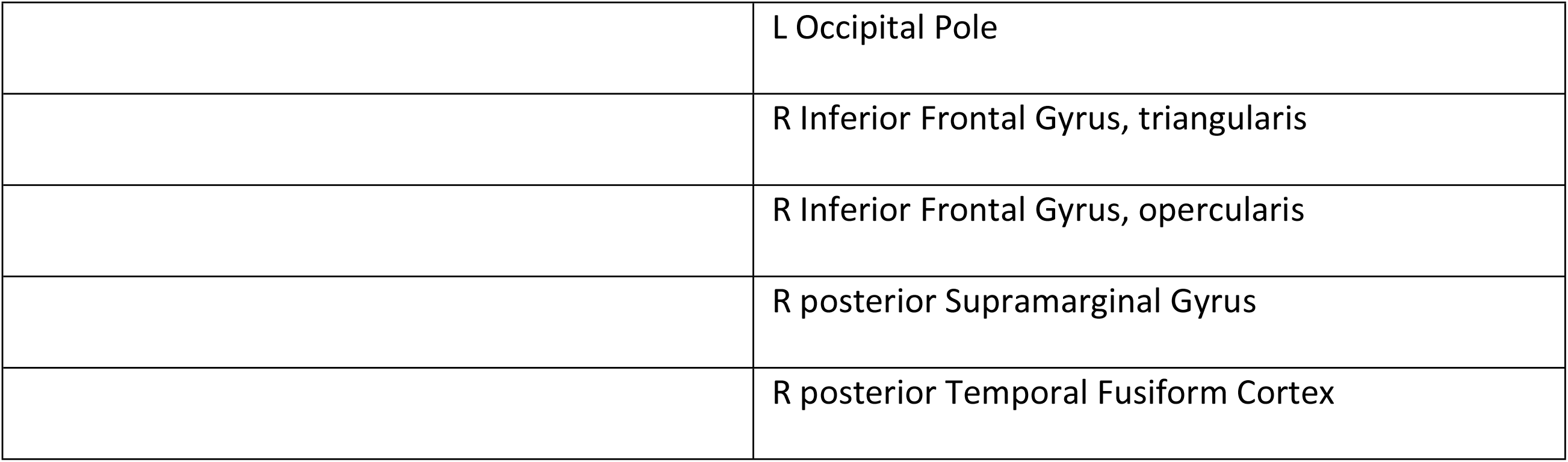
How often do you read fiction? Results of fMRI network analysis. Regions which had a statistically significant number of functional connections with other regions, depending on participant’s score on the question ‘How often do you read fiction?’. Displayed are the labels of the regions (as taken from the Harvard-Oxford atlas) and the number of connections per region (node degree, ordered from largest to smallest). For an illustration of the location of the regions see Figure 1A, and Supplementary Table S1. For an overview of the regions to which the regions in this table were connected to, see Supplementary Materials S3.

Another set of areas was found to have higher node degree depending on the score on the ***ART*** (Table 4, Fig. 5, Figure 6B). This network partially overlapped with the regions showing increased connectivity depending on how much the participant reported to read and included (among others) the bilateral inferior frontal sulci, the anterior medial prefrontal cortex (MPFC),Left angular gyrus, and left temporal pole (see Table 4, Fig. 5, Figure 6B).

**Table 4:**
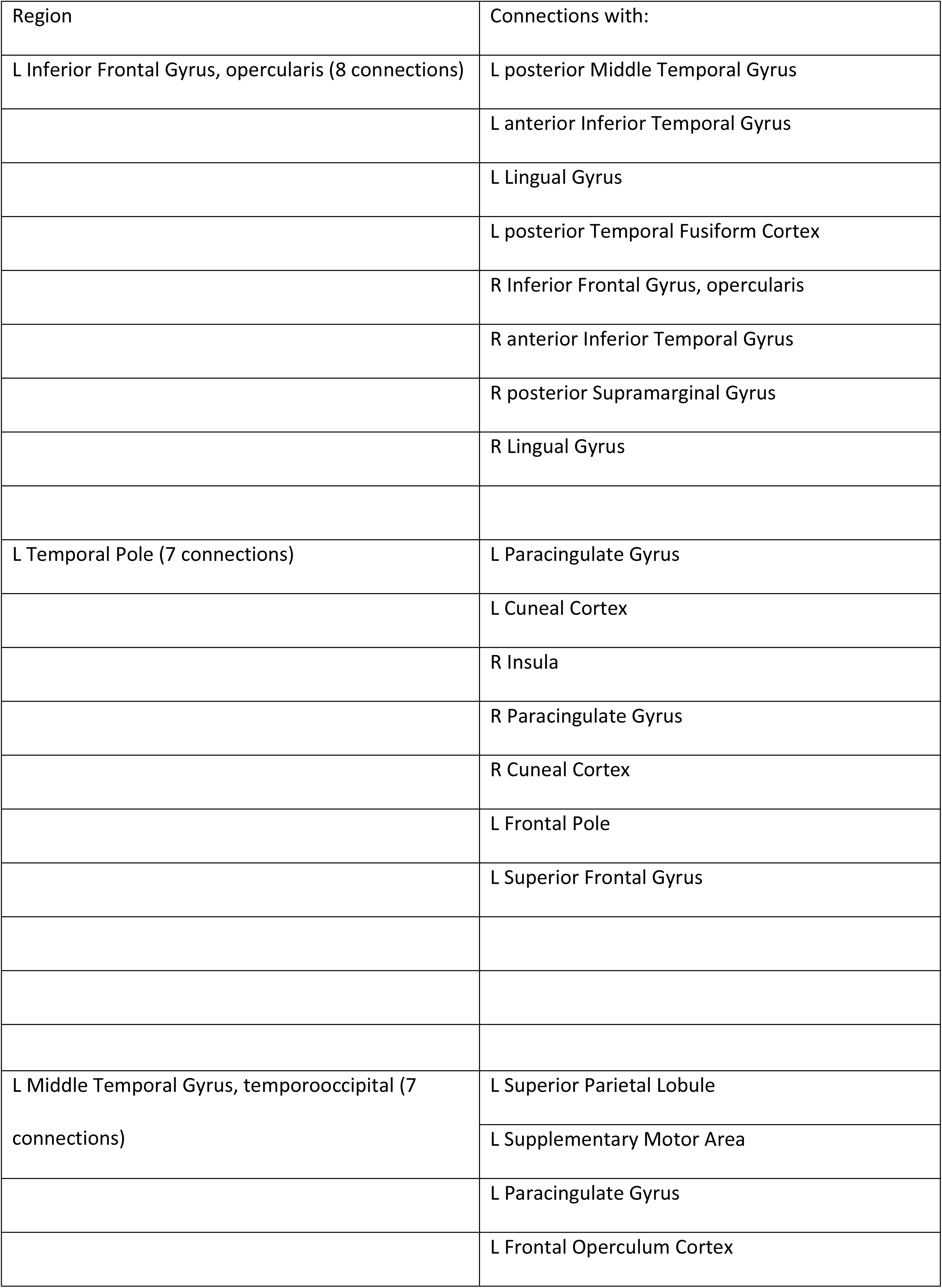
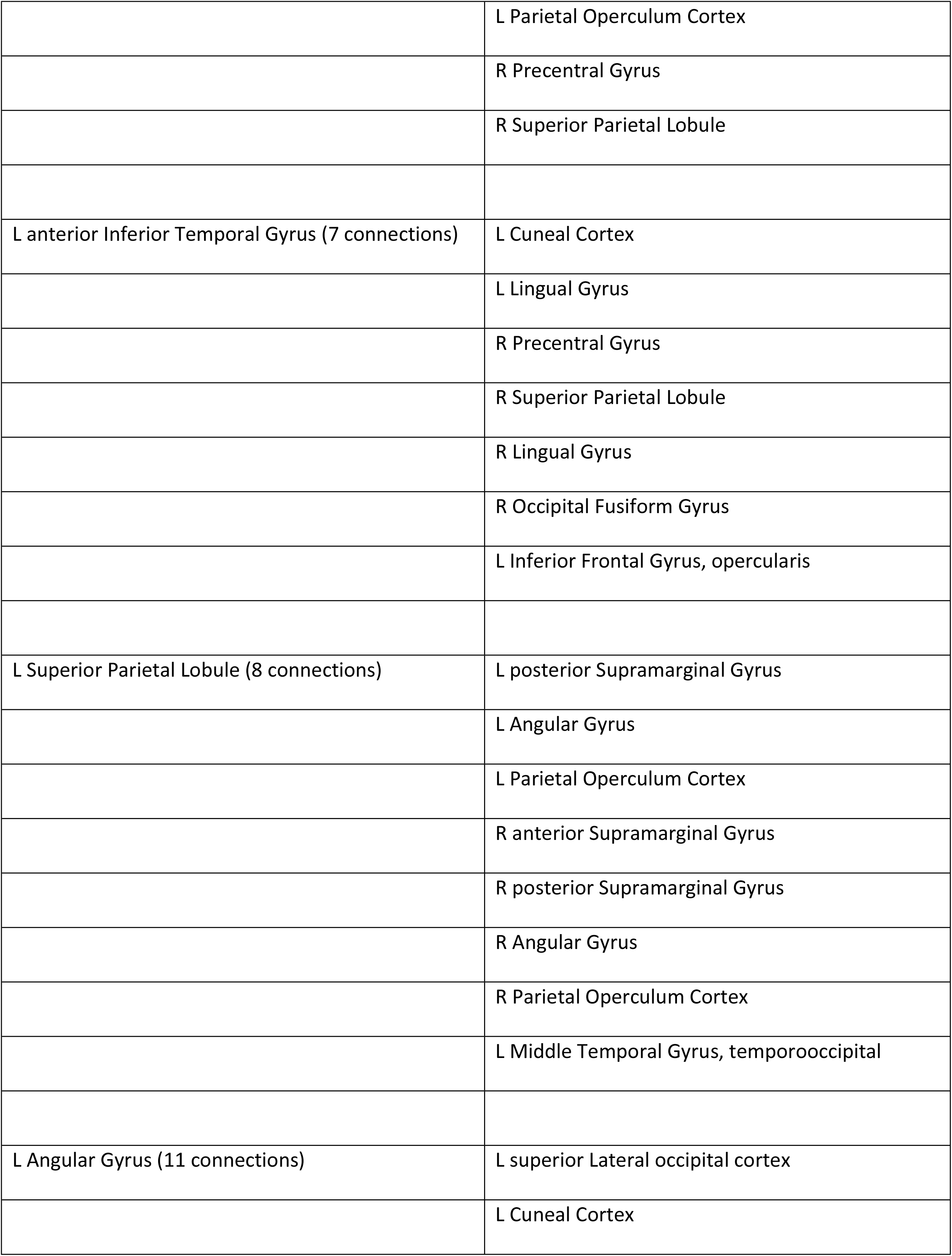
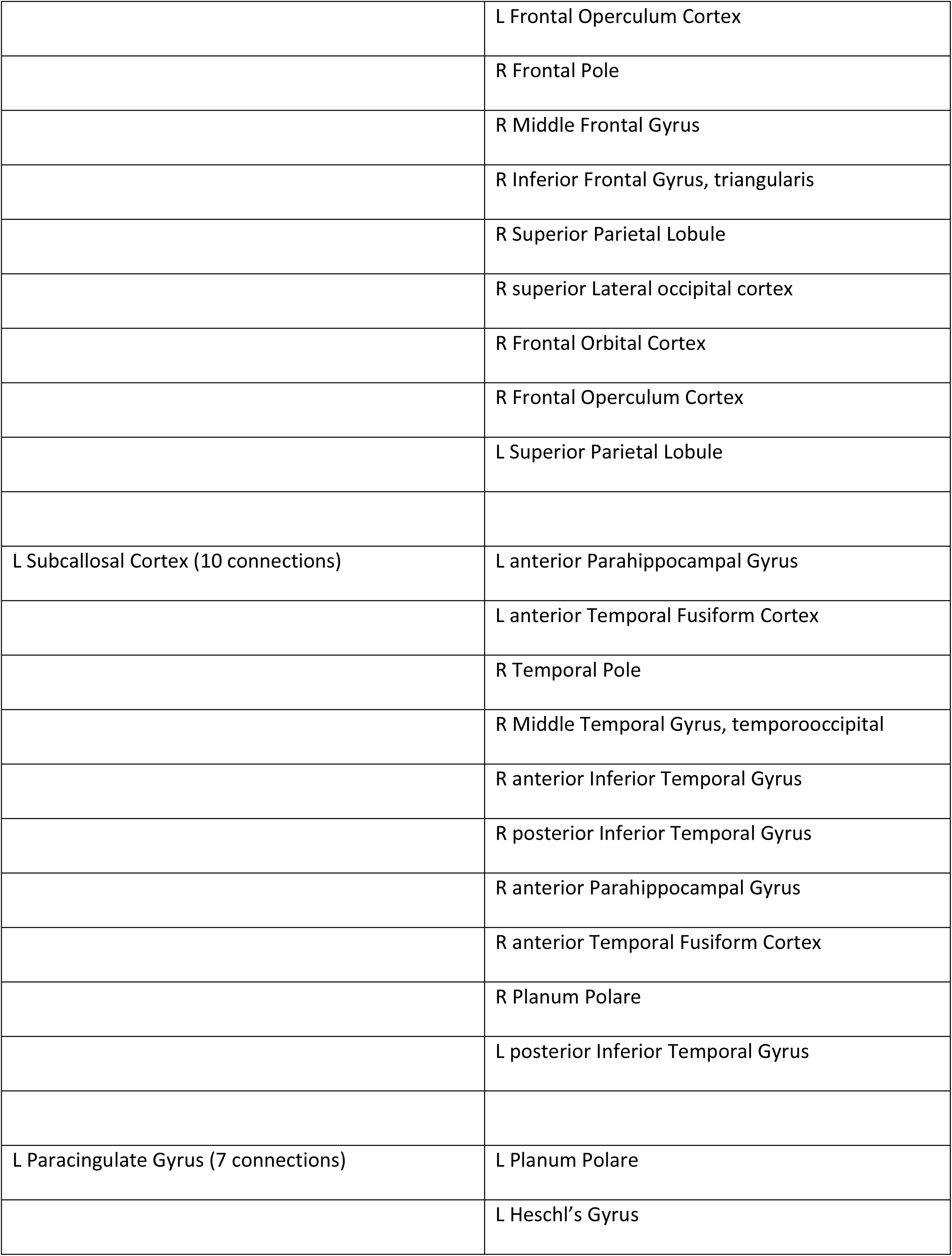
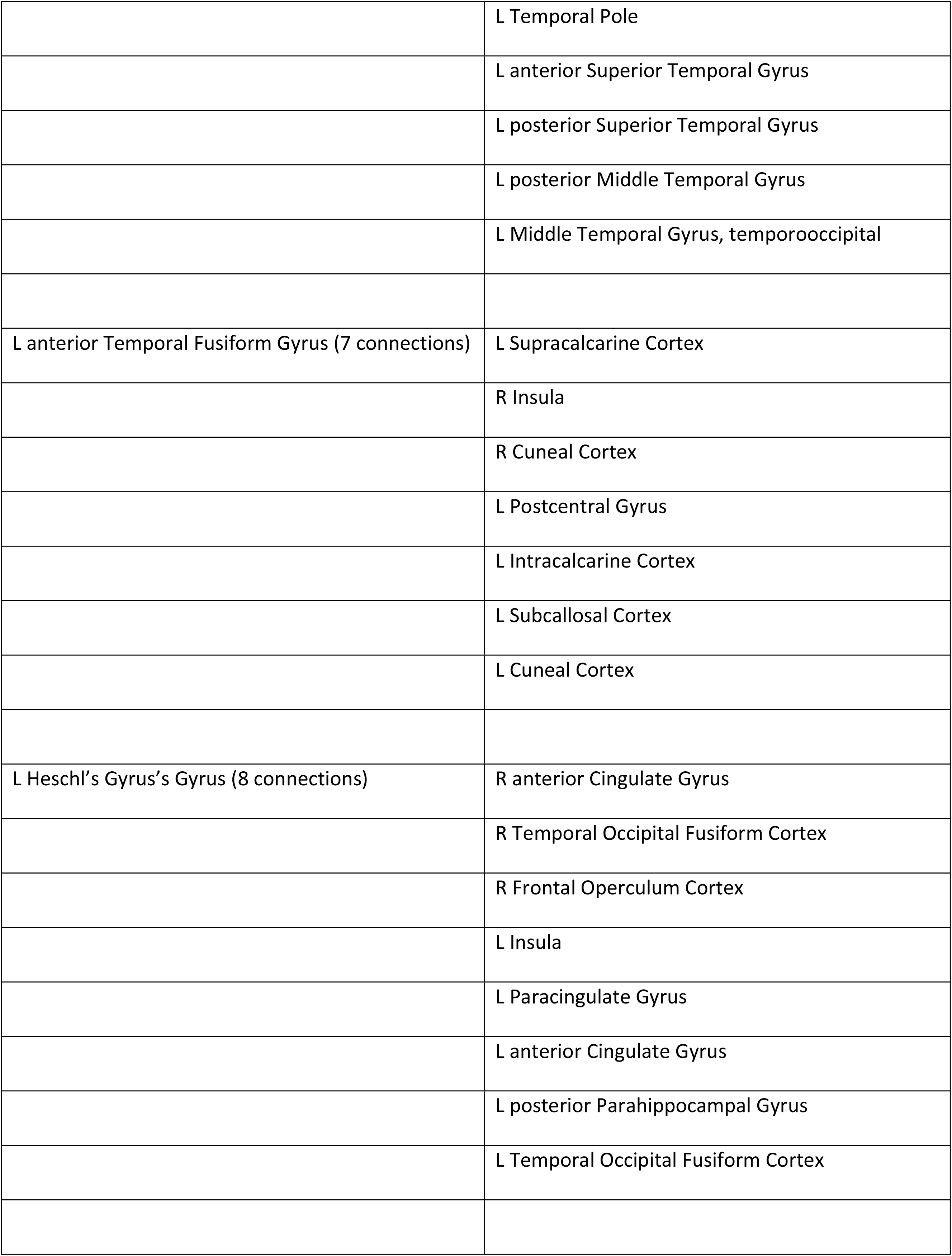
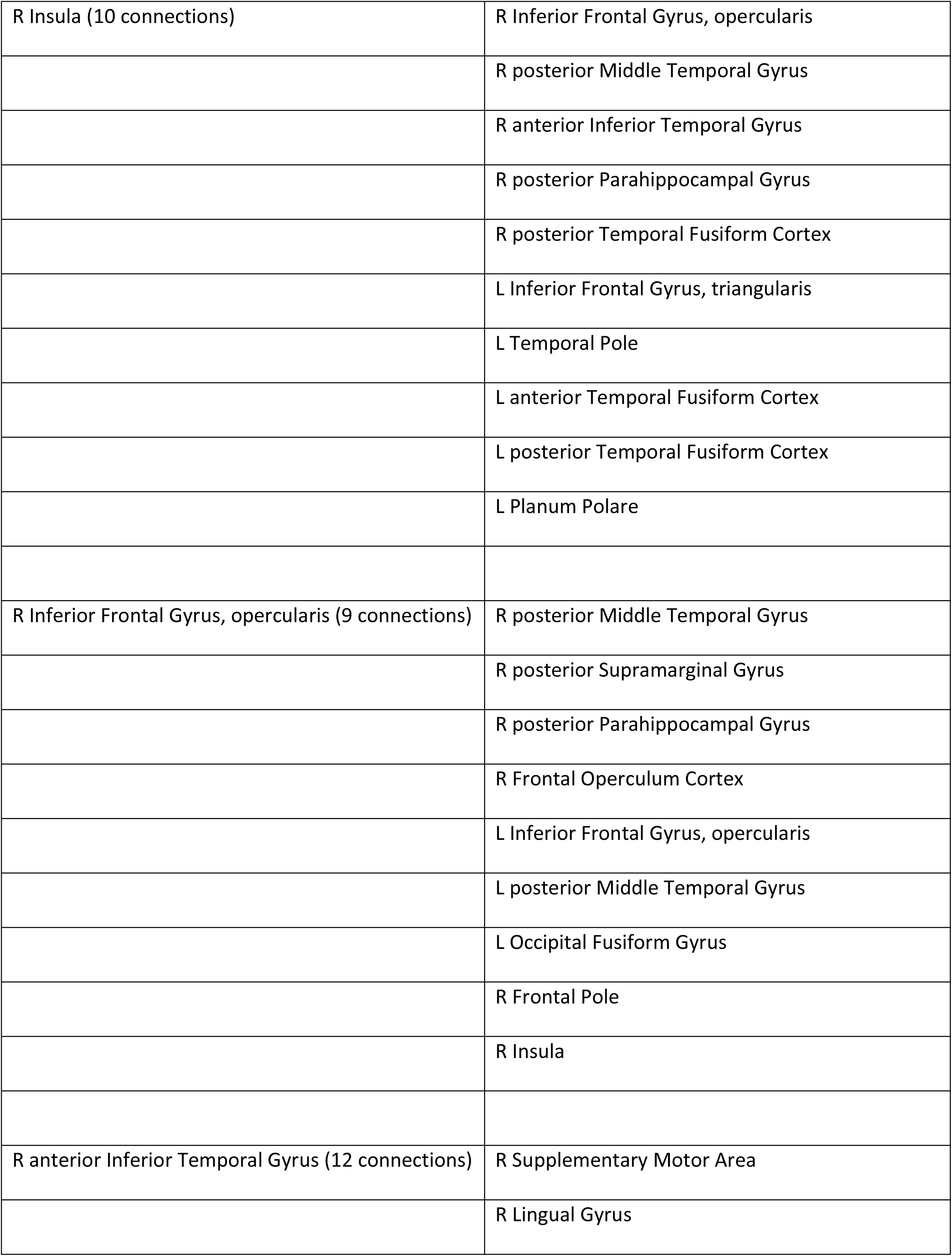
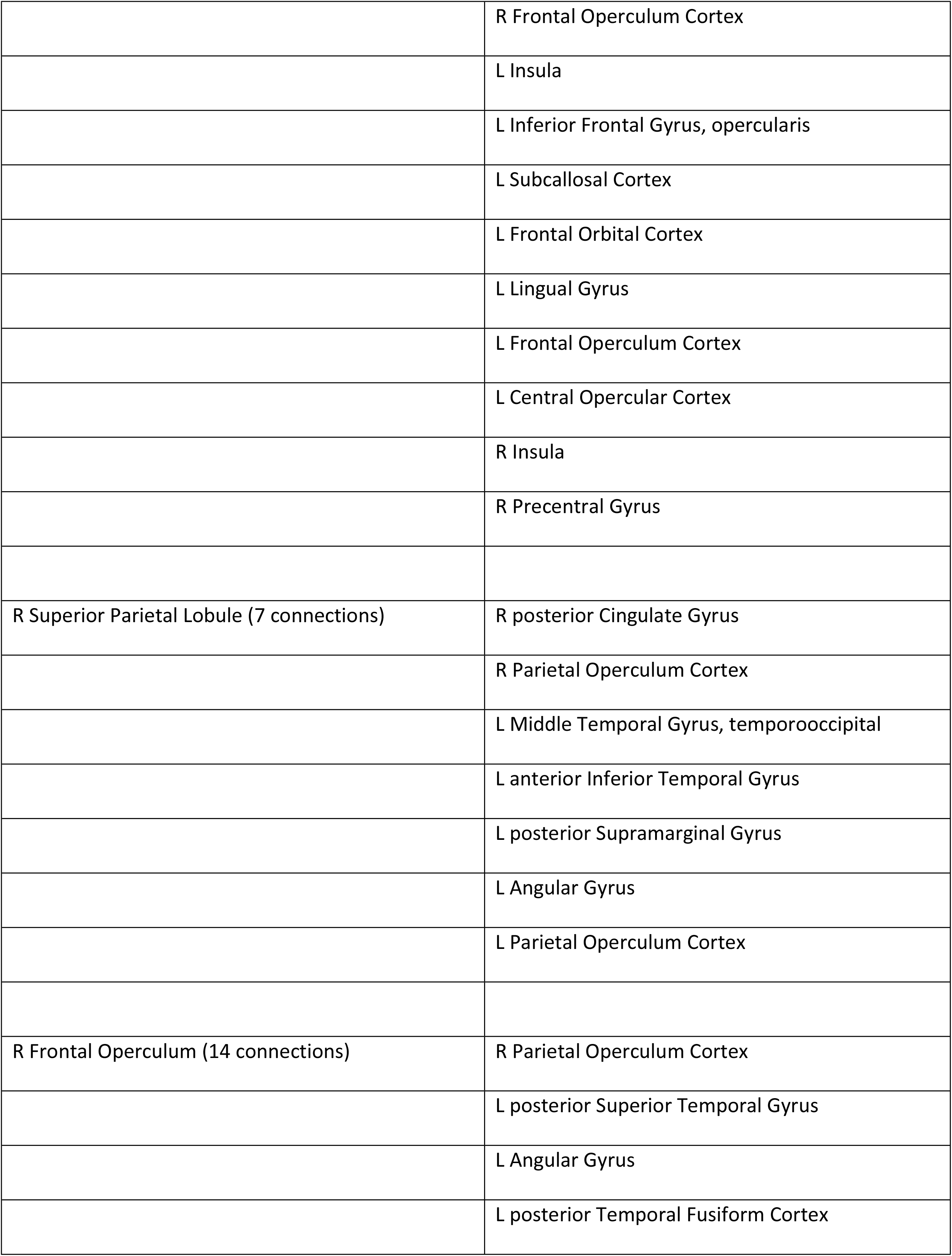
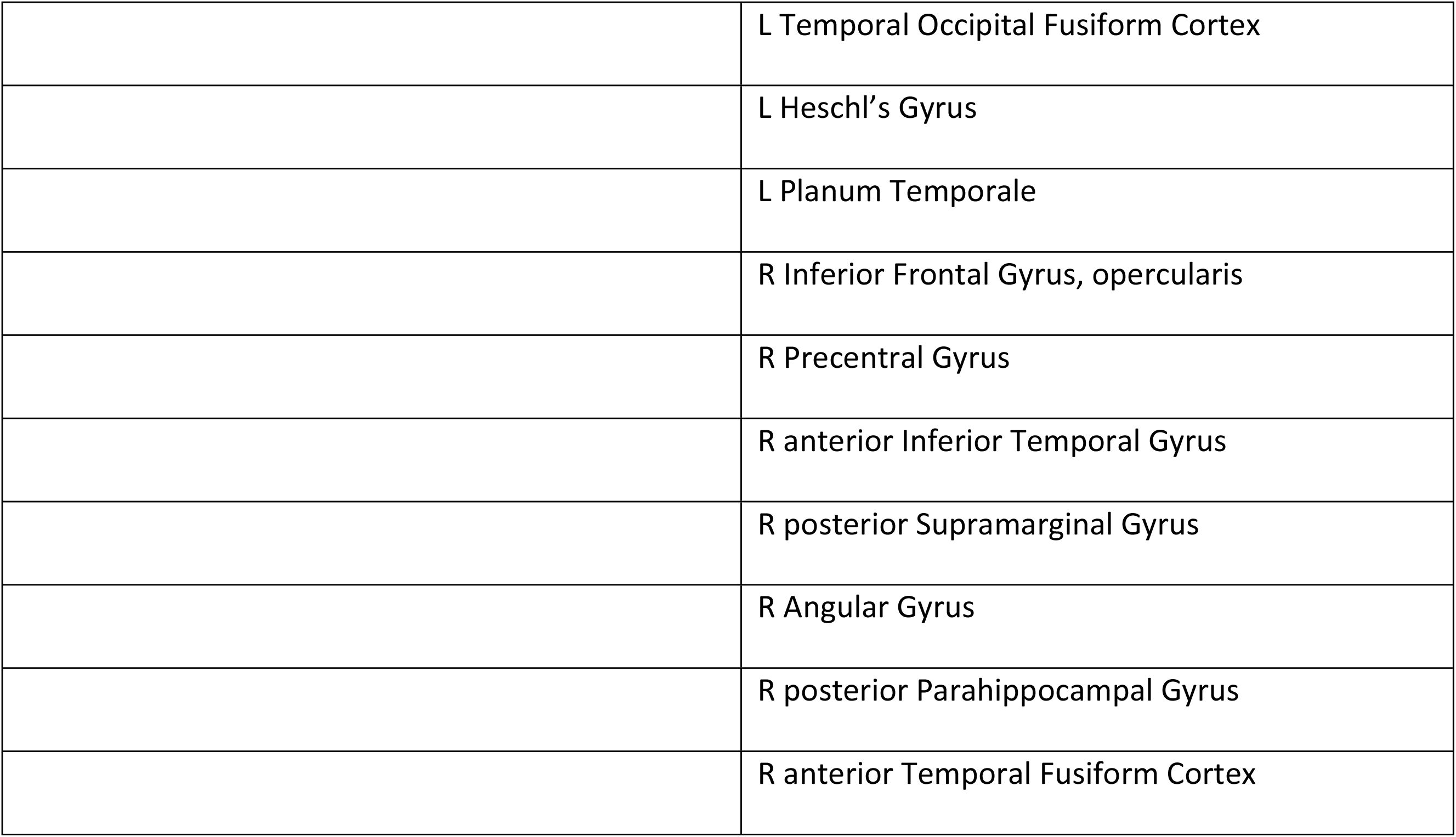
Author Recognition Test (ART). Results of fMRI network analysis. Regions which had a statistically significant number of functional connections with other Regions, depending on participant’s score on the Author Recognition Test (an implicit measure of print exposure, see text). Displayed are the Labels of the Regions (as taken from the Harvard-Oxford atlas) and the number of connections per Region (node degree, ordered from Largest to smallest). For an illustration of the Location of the Regions see Figure 1A, and Supplementary Materials S2. For an overview of the Regions to which the Regions in this table were connected to, see Supplementary Materials S3.

| Region | Connections with: |
| --- | --- |
| L Inferior Frontal Gyrus, opercularis (8 connections) | L posterior Middle Temporal Gyrus |
|  | L anterior Inferior Temporal Gyrus |
|  | L Lingual Gyrus |
|  | L posterior Temporal Fusiform Cortex |
|  | R Inferior Frontal Gyrus, opercularis |
|  | R anterior Inferior Temporal Gyrus |
|  | R posterior Supramarginal Gyrus |
|  | R Lingual Gyrus |
| L Temporal Pole (7 connections) | L Paracingulate Gyrus |
|  | L Cuneal Cortex |
|  | R Insula |
|  | R Paracingulate Gyrus |
|  | R Cuneal Cortex |
|  | L Frontal Pole |
|  | L Superior Frontal Gyrus |
| L Middle Temporal Gyrus, temporooccipital (7 connections) | L Superior Parietal Lobule |
|  | L Supplementary Motor Area |
|  | L Paracingulate Gyrus |
|  | L Frontal Operculum Cortex |
|  | L Parietal Operculum Cortex |
|  | R Precentral Gyrus |
|  | R Superior Parietal Lobule |
| L anterior Inferior Temporal Gyrus (7 connections) | L Cuneal Cortex |
|  | L Lingual Gyrus |
|  | R Precentral Gyrus |
|  | R Superior Parietal Lobule |
|  | R Lingual Gyrus |
|  | R Occipital Fusiform Gyrus |
|  | L Inferior Frontal Gyrus, opercularis |
| L Superior Parietal Lobule (8 connections) | L posterior Supramarginal Gyrus |
|  | L Angular Gyrus |
|  | L Parietal Operculum Cortex |
|  | R anterior Supramarginal Gyrus |
|  | R posterior Supramarginal Gyrus |
|  | R Angular Gyrus |
|  | R Parietal Operculum Cortex |
|  | L Middle Temporal Gyrus, temporooccipital |
| L Angular Gyrus (11 connections) | L superior Lateral occipital cortex |
|  | L Cuneal Cortex |
|  | L Frontal Operculum Cortex |
|  | R Frontal Pole |
|  | R Middle Frontal Gyrus |
|  | R Inferior Frontal Gyrus, triangularis |
|  | R Superior Parietal Lobule |
|  | R superior Lateral occipital cortex |
|  | R Frontal Orbital Cortex |
|  | R Frontal Operculum Cortex |
|  | L Superior Parietal Lobule |
| L Subcallosal Cortex (10 connections) | L anterior Parahippocampal Gyrus |
|  | L anterior Temporal Fusiform Cortex |
|  | R Temporal Pole |
|  | R Middle Temporal Gyrus, temporooccipital |
|  | R anterior Inferior Temporal Gyrus |
|  | R posterior Inferior Temporal Gyrus |
|  | R anterior Parahippocampal Gyrus |
|  | R anterior Temporal Fusiform Cortex |
|  | R Planum Polare |
|  | L posterior Inferior Temporal Gyrus |
| L Paracingulate Gyrus (7 connections) | L Planum Polare |
|  | L Heschl's Gyrus |
|  | L Temporal Pole |
|  | L anterior Superior Temporal Gyrus |
|  | L posterior Superior Temporal Gyrus |
|  | L posterior Middle Temporal Gyrus |
|  | L Middle Temporal Gyrus, temporooccipital |
| L anterior Temporal Fusiform Gyrus (7 connections) | L Supracalcarine Cortex |
|  | R Insula |
|  | R Cuneal Cortex |
|  | L Postcentral Gyrus |
|  | L Intracalcarine Cortex |
|  | L Subcallosal Cortex |
|  | L Cuneal Cortex |
| L Heschl's Gyrus's Gyrus (8 connections) | R anterior Cingulate Gyrus |
|  | R Temporal Occipital Fusiform Cortex |
|  | R Frontal Operculum Cortex |
|  | L Insula |
|  | L Paracingulate Gyrus |
|  | L anterior Cingulate Gyrus |
|  | L posterior Parahippocampal Gyrus |
|  | L Temporal Occipital Fusiform Cortex |
| R Insula (10 connections) | R Inferior Frontal Gyrus, opercularis |
|  | R posterior Middle Temporal Gyrus |
|  | R anterior Inferior Temporal Gyrus |
|  | R posterior Parahippocampal Gyrus |
|  | R posterior Temporal Fusiform Cortex |
|  | L Inferior Frontal Gyrus, triangularis |
|  | L Temporal Pole |
|  | L anterior Temporal Fusiform Cortex |
|  | L posterior Temporal Fusiform Cortex |
|  | L Planum Polare |
| R Inferior Frontal Gyrus, opercularis (9 connections) | R posterior Middle Temporal Gyrus |
|  | R posterior Supramarginal Gyrus |
|  | R posterior Parahippocampal Gyrus |
|  | R Frontal Operculum Cortex |
|  | L Inferior Frontal Gyrus, opercularis |
|  | L posterior Middle Temporal Gyrus |
|  | L Occipital Fusiform Gyrus |
|  | R Frontal Pole |
|  | R Insula |
| R anterior Inferior Temporal Gyrus (12 connections) | R Supplementary Motor Area |
|  | R Lingual Gyrus |
|  | R Frontal Operculum Cortex |
|  | L Insula |
|  | L Inferior Frontal Gyrus, opercularis |
|  | L Subcallosal Cortex |
|  | L Frontal Orbital Cortex |
|  | L Lingual Gyrus |
|  | L Frontal Operculum Cortex |
|  | L Central Opercular Cortex |
|  | R Insula |
|  | R Precentral Gyrus |
| R Superior Parietal Lobule (7 connections) | R posterior Cingulate Gyrus |
|  | R Parietal Operculum Cortex |
|  | L Middle Temporal Gyrus, temporooccipital |
|  | L anterior Inferior Temporal Gyrus |
|  | L posterior Supramarginal Gyrus |
|  | L Angular Gyrus |
|  | L Parietal Operculum Cortex |
| R Frontal Operculum (14 connections) | R Parietal Operculum Cortex |
|  | L posterior Superior Temporal Gyrus |
|  | L Angular Gyrus |
|  | L posterior Temporal Fusiform Cortex |
|  | L Temporal Occipital Fusiform Cortex |
|  | L Heschl's Gyrus |
|  | L Planum Temporale |
|  | R Inferior Frontal Gyrus, opercularis |
|  | R Precentral Gyrus |
|  | R anterior Inferior Temporal Gyrus |
|  | R posterior Supramarginal Gyrus |
|  | R Angular Gyrus |
|  | R posterior Parahippocampal Gyrus |
|  | R anterior Temporal Fusiform Cortex |

No areas survived the statistical threshold as set by the randomization approach for correlations of node degree with the questions ‘***How much do you like reading fiction?***’ and ‘***How many novels did you read last year?***’. There were, however, sizeable correlations between node degree of the regions in the network related to the questions ‘How often do you read fiction?’ (discussed above) and ‘How much do you like reading fiction?’ (rho=0.46, p<0.001), and ‘How often do you read fiction?’ and ‘How many novels did you read last year?’ (rho=0.37, p<0.001). This suggests that the fact that we did not observe networks to the latter two measures (‘How much do you like reading fiction?’, and ‘How many novels did you read last year?’) is due to the statistical thresholding applied.

Given the special interest in the influence of amount of fiction reading on neural connectivity we selected the regions with the highest node degree to the question ‘How often do you read fiction?’ (5 highest degree areas illustrated in Fig. 7, see Fig. 8 for a correlation matrix), and illustrate to which regions they are connected (Fig. 7). The areas with highest node degree are the right inferior frontal cortex (pars triangularis, 26 connections), the posterior part of the right supramarginal gyrus (16 connections), the right middle frontal gyrus (16 connections),Right inferior frontal cortex (pars opercularis, 14 connections), and the left lingual gyrus (13 connections). In figure 7, we see for each of these hub regions (pink regions in subpanels of the figure) the connection strength to other regions which was positively moderated by amount of fiction reading (see Table 3 for a full listing of all regions with high node degree and their connections). As can be seen in the figure, these areas are often connected to each other, with other noticeable connections to the right temporal pole, anterior medial prefrontal cortex, and the precuneus (see also Fig. 8).

**Figure 7:**
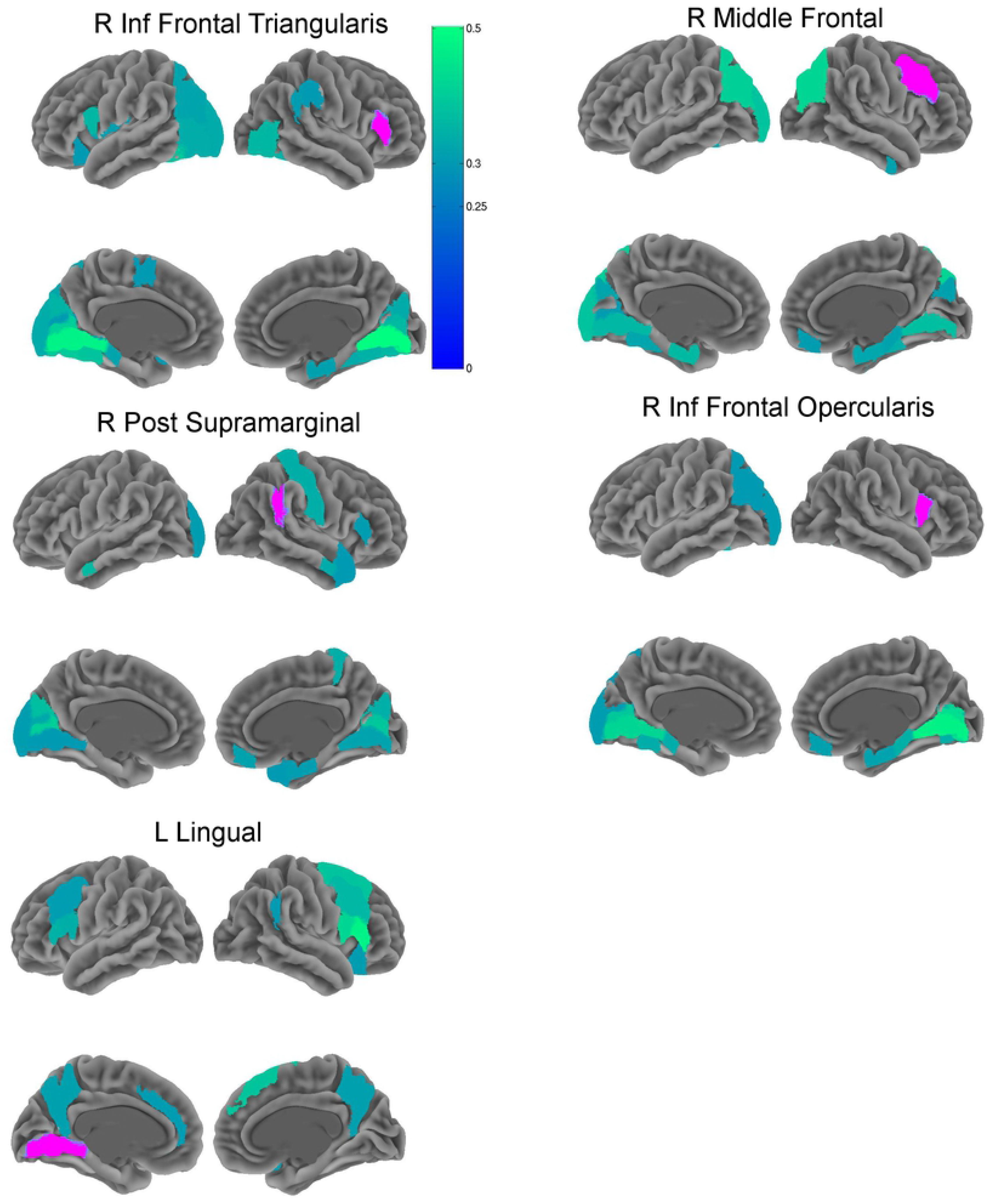
Areas with highest node degree in the analysis to the question ‘How often do you read fiction?’. The five areas which had the highest node degree (for the full set of areas see Fig. 3A and Table 2) are displayed in each sub panel in pink. In blue-green are the areas which had connections to the region, at the threshold which was set for the correlation analysis. Color coding indicates Spearman’s correlation.

**Figure 8:**
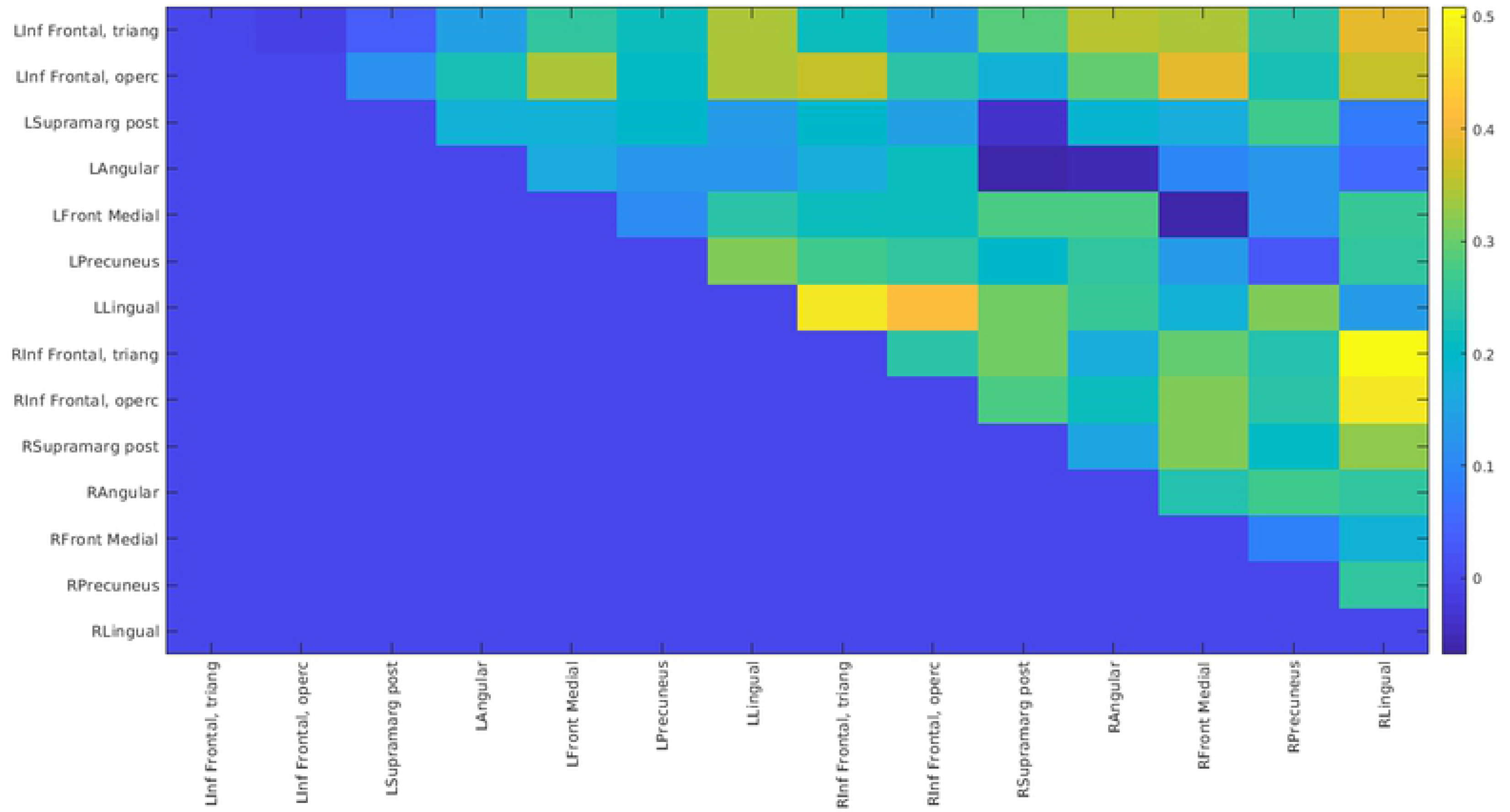
Increased connectivity between regions with high node degree dependent on the question ‘how often do you read fiction?’.

## Discussion

We explored a possible neurobiological basis for the long-standing hypothesis that reading fiction improves social cognition abilities. We tested whether there are differences in connection strengths between cortical regions while people listen to literary narratives depending on how much fiction they read. Because reading fiction is proposed to be a training mode for social abilities, we expected a positive link between reading fiction and connectivity between regions implicated in understanding the intentions, beliefs and feelings of others. The results show that several brain regions that are linked to social cognition and language processing show an increased number of connections with other regions, the higher a participant scored on the question ‘How often do you read fiction?’ (Fig. 3A). The key regions showing increased connectivity in avid readers are the inferior frontal cortices bilaterally, the lingual gyri bilaterally, the right middle frontal gyrus, the posterior part of the right supramarginal gyrus, and the anterior part of the medial frontal cortex. Several of these regions are not only parts of the mentalizing and language networks but also part of the DMN. The DMN seems to be involved in aspects of cognition that are linked to aesthetic experience, self-relevance and self-referential thinking, episodic thinking (28), and non-directed attention (e.g. mind wandering, flow). It is thus not surprising that several studies found it to be involved in narrative processing (6,25,26). A fundamental aspect of engaging with literature is the aesthetic appraisal of the language used and the plot built (e.g. suspense and twists), but also the cultural, social, and self-referential search for greater meaning. Narrative processing naturally involves episodic structuring of events in a story. Immersing in a story also involves decrease in executive control and directed attention, as participants allow the story to lead their attention instead of actively seeking information.

Before we discuss the most important regions which showed sensitivity to the amount of fiction reading, an important point needs to be made. In our analysis, we establish which regions have more functional connections with other regions during listening to a literary narrative, depending on how much participants read fiction. The hypothesis (as discussed above) is that this is a training effect: through repeated exposure to fiction, these areas are activated together more frequently, and hence form a network that is more easily activated as compared to people who have less exposure to fiction. While this is a viable hypothesis, an alternative hypothesis is that people with stronger connections in these networks enjoy reading more and hence are more likely to be avid readers. These two hypotheses are not mutually exclusive, and it is important to point out that our experimental design is agnostic to the directionality of this relationship.

All regions that show differences depending on how much people read are regions that are typically found in tasks related to social cognition and language processing, including sensory and motor areas that are typically involved in processing semantic information about events and characters. The left and right inferior frontal regions are the first notable regions whose number of connections showed a sensitivity to amount of fiction reading. The involvement of these regions as part of the language network is well-established (40), and especially right inferior frontal gyrus (RIFG) has been implicated in discourse comprehension (54,55). In the present results the RIFG formed connections with the left inferior frontal cortex, the lingual gyri, and the anterior part of the medial frontal cortex - a prominent region in the mentalizing network (Fig. 7, Fig. 8). This suggests that the RIFG could be an interfacing region between the language network and parts of the mentalizing network.

The region to region connectivity pattern that was most affected by how much participants read was the connectivity of the left and right inferior frontal gyri to the left and right lingual gyri. Left IFG is a prominent region in language processing, linked to semantic selection and coordination, whereas right IFG seems to be crucial for discourse processing (see discussion above). The connectivity profile of the left lingual gyrus (one of the regions with the highest node degree) was such that it connects with inferior frontal cortices bilaterally, the right angular gyrus, as well as the precuneus bilaterally (Fig. 8). Both, the precuneus and right temporo-parietal regions are part of the extended language network (54). The lingual gyrus is not typically considered part of the language network (and hence was not hypothesized to be implicated in this study). However, several fMRI studies report that the lingual gyri are activated during language comprehension (56–58). It has also been shown that the cortical thickness of this region co-varies with lateral temporal, inferior parietal and inferior frontal regions often implicated in language, thus indicating that these regions experience common activation, presumably related to language comprehension (59). A full discussion of this region and its potential role in language comprehension is beyond the scope of this paper (42). It does seem however, that it may be an area which has been somewhat overlooked by the field.

Another notable region which was correlated with the amount of fiction participants read is the anterior part of the medial frontal cortex. This part of the brain has been repeatedly and consistently shown to be involved in mentalizing, and can be considered a core node of the mentalizing system (22,34,35). The sensitivity of the number of connections this region forms during narrative to the amount of fiction that participants read is the strongest evidence from the present data for the hypothesis that engaging with fiction might train social cognition brain networks.

Two other areas which have been previously implicated in discourse comprehension and showed an effect in the present study are the cuneus and precuneus. It has been suggested that these posterior midline structures serve coherence establishment, and / or situation models (54,60). These regions are also often implicated in thinking about the past and the future, hypothetical thinking, and daydreaming (13). Interestingly, some studies suggest that these areas may generally be implicated in the recognition of structure across time, and this function is not restricted to language. For instance, Tobia and colleagues observed activation increases in posterior cingulate cortex, cuneus and precuneus, when participants noted changes in regularity in tone sequences, that is, in non-linguistic stimulus sequences (61).

Two final areas to note are the posterior supramarginal gyri. The right posterior supramarginal area was connected with right inferior frontal regions, several occipital regions, the postcentral sulcus, and the right temporal pole. In a meta-analysis on the neural basis of semantic processing, Binder and colleagues conclude that the angular gyrus - an area close to the posterior supramarginal gyrus - ‘occupies a position at the top of a processing hierarchy underlying concept retrieval and conceptual integration’ (37). In the Harvard-Oxford atlas we used for the present study, the angular gyrus is separated from the posterior supramarginal gyrus (areas 21 and 20 in Fig. 1). We tentatively suggest that there might be some difference in labeling, meaning that what Binder and colleagues labeled as the angular gyrus could functionally be the posterior supramarginal gyrus in the current study. These regions are also adjacent to the temporo-parietal junction, which is (bilaterally) considered central to the ToM network. This spatial proximity might support an interface between cognitive systems for the integration of semantic and social information.

In discussing the results, we have somewhat artificially distinguished between neural areas important for language and neural areas important for social cognition. This distinction is well-founded since language and social cognition are cognitively and neurally distinguishable (23,62) and have historically been investigated by different subfields. At the same time, it should be clear that conveying and understanding intentions, feelings, and beliefs is a primary goal of language use, narrative, and indeed of human communication at large.

It might also seem non-intuitive that a lot of areas showing sensitivity to how much fiction people read are in sensory and motor areas. However, fiction stories are typically about characters, descriptions of the worlds they live in, and what happens to them. From this perspective, it is not surprising that how much people read might also train their ability to imagine or simulate perceptual descriptions or events taking place in the respective fictional worlds in domain specific areas that process sensory or action information.

While the evidence of a relation between lifetime fiction exposure and differences in functional network connectivity during listening to literary stories is purely correlational, our study confirms the link between the neural processes involved in reading and in social cognition. We show that reading fiction is linked to increased number of functional connections within and between language and mentalizing regions and to other parts of the brain. Apart from established language-related regions like inferior frontal areas, regions recognized as the extended language network seem to play an important role in the brain network processing narratives.

Interestingly, we found no correlation between trait empathy (EQ) and reading in our sample. Previous studies also failed to find a link between self-reported trait measures for social skills and reading behavior measures (12), while self-reported happiness in social relationships was found to be related to fiction reading in the same sample. It is possible that trait empathy is in fact not related to reading, not trainable, or that the social cognition skills that might be trained by reading are not the same as measured by the EQ questionnaire. The EQ was originally designed as a diagnostic tool for people with clinically relevant impairments and not to be a sensitive measure to scale individual differences in trait empathy in healthy adults. While the EQ is still a widely used measure for individual differences in social skills, it might in fact be insensitive for measuring behaviorally relevant differences in healthy adults. In this study, we used the EQ as a control analysis to test if our analyses are sensitive to region-region connectivity differences in social cognition networks depending on one individual difference score.

The causal link between exposure to and initial interest in fiction is interesting, but far beyond the scope of this paper. What is important to note is that the link between a life-time exposure to fiction and social cognition seems to be more robust than brief exposure to fiction, as was shown by two recent large-scale studies (14,15) which found that there is no immediate effect of social cognition performance improvement after a short exposure to fiction, but long-term fiction exposure was positively related to social cognition skills.

It is possible that there are other individual differences overlapping with how much people read that could explain the results. For example, IQ is related to language skills that in turn are related to how much people read (36). Since for this study such global individual differences in cognitive skills were not collected, we refrain from speculating about them.

How explicitly people mentalize with fictional characters when engaging with fictional narratives and whether this is similar to mentalizing with other humans remains an open question. A recent study shows overlap between neural areas involved in explicit versus implicit mentalizing tasks (63), suggesting overlapping cognitive resources. It however seems viable that implicit attribution of mental states to a fictional character is not necessarily qualitatively the same as explicitly ascribing intentions or beliefs to another real-world agent, particularly in an interactive setting.

Our results provide evidence for a possible mechanism that could support fiction reading as a training mode for social cognition by facilitating (task based) connectivity between brain regions involved in ToM. At the same time, we show that it is not just social cognition which would be trained by regularly engaging with fiction. Reading fiction is correlated with a network which is important for discourse understanding, which includes, but is not limited to, mentalizing. From our results, it is evident that more general language skills should be also included in the list of abilities that could be trained by reading. This is to be expected given the evidence in the scientific literature for the relation between print exposure and language abilities (36).

Our findings are largely in line with those of Tamir and colleagues who found that activations in parts of the mentalizing network are modulated by amount of fiction reading as implicitly tested in participants (13). It should be stressed that the increased between-region connection strengths that we observed in the present study cannot be explained by reference to resting state fMRI (rs-fMRI). In the present study we effectively canceled the contribution of what happens during the resting state with our baseline reference.

Our results are evidence for a correlational link between brain network interaction in mentalizing and language processing areas of the brain, but the question of causality and directionality is still unanswered. Longitudinal brain connectivity studies are needed to see whether fiction consumption leads to actual changes in brain connectivity patterns to draw causal conclusions. Our study only tested for two short stories that are very similar in the type of suspense and emotion. The generalizability of these findings remains to be tested with more diverse materials. It is further unclear, if there is a difference between quality and quantity of fiction consumed and whether these effects actually depend on verbal narratives or whether comparable links can be confirmed for other types of narratives such as comics, movies, or games.

One important limitation of this study is that we lack a measure of actual performance in a ToM task. One often employed performance measure of social cognition abilities that has been used by previous research on the influence of fiction reading on social cognition is the ‘Reading the Mind in the Eyes’ test (12,14,18). It is a measure of recognition of mental and emotional states from static black and white images of human eyes. While this test is a widely used measure, the ecological validity of recognizing mental states and emotions from static black and white images is questionable (64) and it also has been criticized for poor internal consistency, high dependence on vocabulary and generally there is mixed evidence for convergent validity with other social cognition measures (65).

We also did not have measures of quality of social life participants have. Whether social cognition abilities (and social cognition brain network connectivity) are a reliable predictor of social life quality and satisfaction is an interesting research question that should be addressed in future studies. If the accumulating evidence for the link between fiction reading and social abilities can be confirmed by studies testing the causal link between the two, this might be a promising avenue for potential treatments for social disorders and quality of social life improvement. We further only counted the number of fictional works or books people consume instead of time spent engaging with stories. This might be an important factor that should be tested. In addition, our sample of participants only included young highly educated adults. It would be interesting to see if these effects generalize over the human life span and to communities in which storytelling fulfills different functions than in Western cultures.

While there is the sort of trivial case that language (or its subprocesses) might also be a cognitive ability trained by reading and not only mentalizing, it is also obvious that many other cognitive functions might be involved in (and possibly trained by)Reading fiction. For example, complex levels of causality (causal reasoning), good and bad characters (morality judgements), descriptions of environments (mental imagery and spatial cognition), are often involved in narratives and might potentially train respective neural networks when people engage with stories. Engaging with fictional characters and their emotional worlds might further train the ability to not only mentalize with others but also improve the ability to communicate about emotions and believes in the real world. Future research is needed to explore these interactions and potential applications.

In sum, we show that areas that are part of the mentalizing and language networks are more strongly connected with other regions in people who more regularly read fiction as compared to those who read less. Our study has high ecological validity due to the fact that our participants listened to entire literary stories from beginning to end instead of excerpts or texts created specifically for the purpose of experimental studies usually used in comparable experiments. We hypothesize that our findings regarding the mentalizing network would generalize to other types of fiction (e.g. movies, theatre, comics), whereas the influence on the language network may be more confined to actual *reading* of or listening to fiction. This is an avenue that future studies should explore. Another important issue for future research is whether and how the link between fiction reading and social cognition can inform actual behavior in the real world. Some studies have used prosocial behavior as a dependent measure in studying the effect of literary reading (16), and it would be important to better understand how actual behavior is influenced by engaging with fiction (66). Most importantly, our study illuminates why engaging with stories can serve such important personal and cultural functions by providing evidence for the potential underlying neural mechanism by which one cognitive function can be ‘trained’ by another by synchronizing networks of brain regions during simultaneous activation.

## Author contributions

The data was collected and preprocessed by FH under supervision ofRW.RW analyzed the data. Both authors wrote this manuscript.

## Acknowledgements

We thank Uri Hasson for pointing us to the work on the lingual gyrus. Paul Gaalman and Jeff Martin helped with data acquisition.

